# Dissecting the effects of single amino acid substitutions in SARS-CoV2 Mpro

**DOI:** 10.1101/2025.03.14.643358

**Authors:** Shwetha Sreenivasan, Joseph D. Fontes, Liskin Swint-Kruse

## Abstract

Successfully predicting the effects of amino acid substitutions on protein function and stability remains challenging. Recent efforts to improve computational models have included training and validation on high-throughput experimental datasets, such as those generated by deep mutational scanning (DMS) approaches. However, DMS signals typically conflate a substitution’s effects on protein function with those on *in vivo* protein abundance; this limits the resolution of mechanistic insights that can be gleaned from DMS data. Distinguishing functional changes from abundance-related effects is particularly important for substitutions that exhibit intermediate outcomes (*e.g.,* partial loss-of-function), which are difficult to predict. Here, we explored changes in *in vivo* abundance for substitutions at representative positions in the SARS-CoV-2 Main Protease (Mpro). For this study, we used previously published DMS results to identify “rheostat” positions, which are defined by having substitutions that sample a broad range of intermediate outcomes. We generated 10 substitutions at each of six positions and separately measured effects on function and abundance. Results revealed an ∼45-fold range of change for abundance, demonstrating that it can make significant contributions to DMS outcomes. Moreover, the six tested positions showed diverse substitution sensitivities for function and abundance. Some positions influenced only one parameter. Others exhibited rheostatic effects on both parameters, which to our knowledge, provides the first example of such behavior. Since effects on function and abundance may arise through different biophysical bases, these results underscore the need for datasets that independently measure these parameters in order to build predictors with enhanced mechanistic insights.

**ABSTRACT for broader audience:** Changing one amino acid in a protein can affect its function, its abundance, or both. Understanding these separate effects will help scientists predict how protein changes alter biology, which is important for understanding pathogen evolution, improving personalized medicine, and bioengineering. This work reports a study to experimentally separate these effects for the main protease of SARS-CoV-2 and suggests strategies for building better prediction models for emerging variants of this key viral protein.

## INTRODUCTION

Accurately predicting the outcomes of single amino acid substitutions is essential for advancing protein evolution, personalized medicine, and bioengineering.^1; 2^ However, despite extensive efforts over three decades, advances in computational predictions have generally been incremental.^3–6^ This indicates that the fundamental biochemical and biophysical mechanisms underlying the protein sequence-structure-function relationship – which form the bases of algorithmic assumptions – are not yet fully understood.

Understanding these mechanisms is particularly challenging for substitution outcomes that are not binary. Instead of leading to complete loss or retention of wild-type (WT) function, many substitutions result in intermediate or enhanced outcomes. Such outcomes are prevalent at “rheostat” positions, where the set of possible substitutions results in a graded functional change (*e.g.,* **Supplementary figure 1** and **Supplementary table 1**).^7–9^ Indeed, predictions for substitution outcomes at rheostat positions are worse than predictions for other types of positions.^9^

To address these challenges, the field has recognized that training and validating computational models requires datasets that are not biased by selections for specific substitutions or positions, to capture the full spectrum of possible outcomes. However, experimentally testing all 19 substitutions at every position in a protein using conventional biochemical assays is challenging. To circumvent this, high-throughput techniques like deep mutational scanning (DMS)^10–12^ have emerged as tools that enable the systematic assessment of substitution effects on proteins.

In DMS, the first step is to create a large library of protein substitution variants via site saturating mutagenesis^13^ across the full-length protein or a region/domain of interest.^14;15^ In a commonly-used DMS approach, the substitution libraries are then introduced into a biological system such as yeast,^16^ bacteria,^17^ or mammalian cells^18^. The cells are then placed under selective conditions, such as drug exposure, and screened for a desired cellular phenotype or reporter outcome.^19; 20^ Finally, high-throughput sequencing techniques^21; 22^ are used to identify substitutions that impacted each cell’s fitness (and thereby infer the effects on the protein) (*e.g.,* ^23^) or *in vivo* activity of the reporter (*e.g.,* ^24; 25^). Regardless of experimental design, measured outputs are often referred to as “functional scores”.^26^

Several factors complicate interpreting the scores from high-throughput DMS assays.^8; 27^ First, different choices of selection pressures can change the fitness/growth measured for substitution variants^28^, and none of these pressures may actually represent physiological conditions.^27^ Second, many studies assume that fitness or growth levels directly correlate with protein function.^29^ For example, variants with worse fitness are interpreted as having worse function, whereas those with enhanced function (beyond that of the wild-type variant) exhibit improved function.^10^ However, a fitness or growth advantage might not be directly proportional to a protein’s functional changes.^26; 30^ Finally, in many DMS studies, the measured outcomes reflect the net effect of each substitution. That is, changes in the observed “functional scores” could arise from affecting any of several protein parameters^31^, including solubility,^32^ stability,^33; 34^ binding affinity,^35–37^ allosteric regulation,^38–40^ ligand specificity^41; 42^, folding rate^43^, translation rate,^44^ and/or enzyme activity^38; 45; 46^. Thus, the outcomes monitored by DMS cannot be assigned to individual aspects of a protein’s function.

The use of “combined” signals presents a new challenge for training and validating computational predictors: Individual protein parameters can be altered through distinct biophysical processes, and each parameter may necessitate specialized predictive models that must then be integrated to generate an overall prediction model. Thus, constructing such models requires experimental datasets that, at the very least, capture changes in both function *and in vivo* abundance. To date, only a limited number of DMS studies report both parameters.^18; 47; 48^ Instead, many studies explicitly assume that single amino acid substitutions do not alter *in vivo* abundance, equating overall growth or fitness changes solely to functional changes.

The use of “combined effects” for building computational predictors may be especially problematic for substitutions with intermediate outcomes. To explore this, we independently measured the effects of single amino acid substitutions on function and *in vivo* abundance in the SARS-CoV-2 main protease (Mpro). Our studies included several positions that showed a high frequency of intermediate outcomes in a published DMS study.^49^ Strikingly, despite sampling only 60 of the 5814 possible Mpro substitutions, this set captured a ∼45-fold range of change in *in vivo* abundance. Moreover, among the six positions tested, we observed diverse combinations of substitution sensitivities for both function and abundance, highlighting the complexity of substitution effects that can arise from single substitutions. To our knowledge, this work provides the first examples of rheostat positions that control both protein function and *in vivo* abundance. As such, these results highlight another complex outcome – the simultaneous tuning of multiple protein parameters – that can arise from substitutions at rheostat positions.

## RESULTS

### In prior DMS studies, Mpro positions exhibited diverse substitution sensitivities

Mpro, also known as the 3C-like protease or Nsp5,^50^ is a highly conserved cysteine protease essential for the replication of coronaviruses^51^ and the target for nirmatrelvir, an FDA-approved treatment for COVID19^52^. Structurally, Mpro exists as a homodimer, and dimerization is essential for its proteolytic activity.^51^ Given Mpro’s role in SARS-CoV-2 replication and the number of changes observed over the course of the pandemic (with single amino acid substitutions observed for at least 287 of 306 positions)^53; 54^, it is useful to understand how single amino acid substitutions alter its function and *in vivo* abundance.

In two previously published DMS studies^49; 55^, the authors created libraries containing >5000 single amino acid substitutions in Mpro. The goal of the first study (in 2022) was to identify positions where most substitutions abolished its activity, with the rationale that they might be good drug targets. The goal of the second study (in 2024) was to detect hyperactive Mpro substitutions, to assess whether their enhanced outcomes enabled drug resistance. In these contexts, abundance changes were perhaps unimportant and did not affect the interpretations made by the authors. Nevertheless, a comparison of the two studies suggests that changes in abundance do affect the assay results.

In the 2022 study^49^, each substitution’s phenotype was assessed using three different types of output: cleavage of a transcription factor that controlled expression of a reporter protein, cleavage of a substrate linked to a fluorescent reporter (FRET), and cleavage of endogenous host proteins that altered yeast growth. Results were assessed using deep sequencing to measure the enrichment or depletion of each substitution in the library, which was then interpreted as the substitutions’ effects on Mpro protease activity. Results were generally in good agreement with each other **(Supplementary figure 2a-c)**. No substitutions with enhanced activity (greater than WT) were detected. The greatest number of positions with intermediate activities were detected using the growth assay^49^, so we used this data set as a reference for our current work (**Figure 1** and **Supplementary figure 2)**.

**Figure 1.**
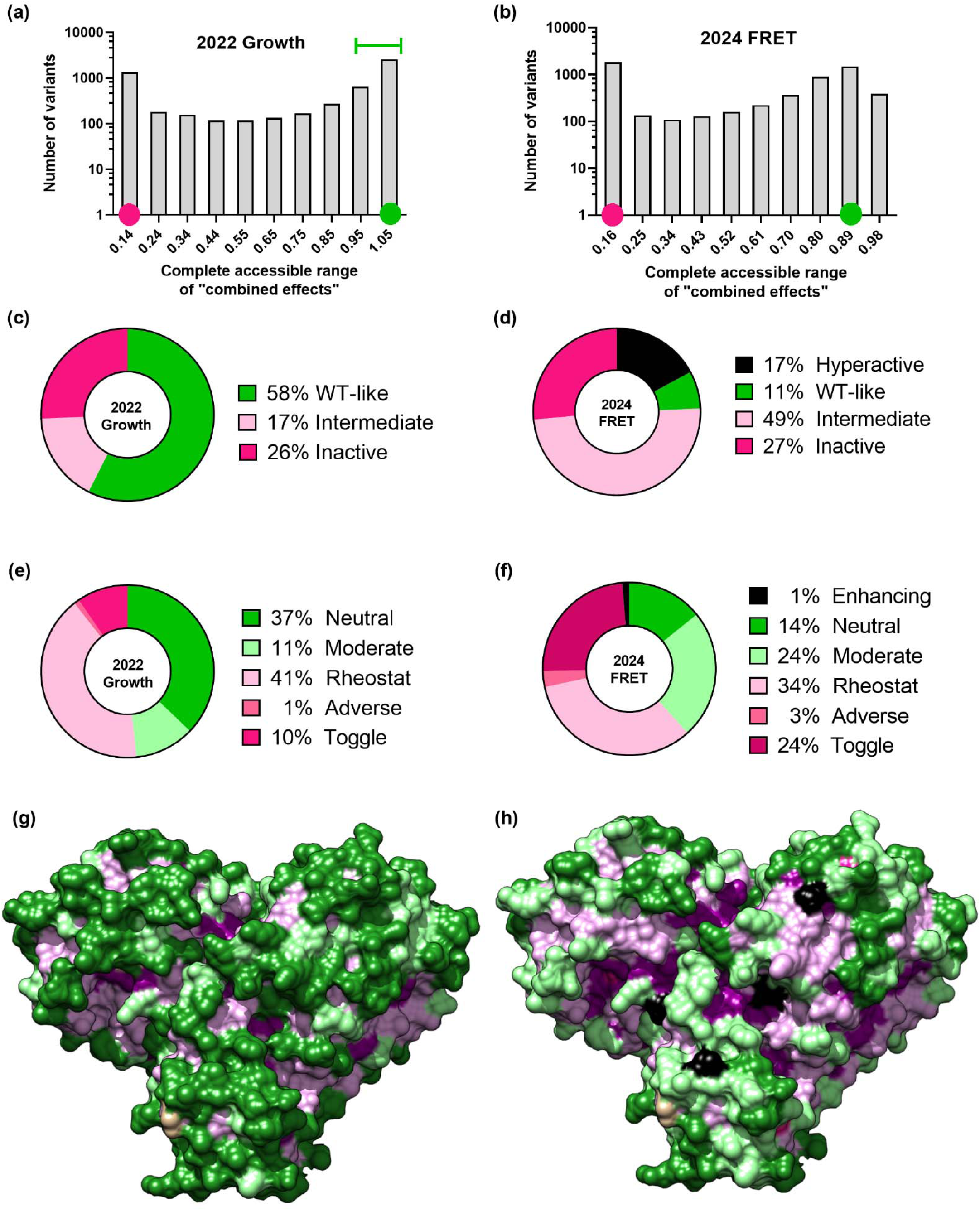
Comparative classification of Mpro substitution sensitivities using datasets from Flynn et al., 2022^49^ and Flynn et al., 2024^55^. **(a, b)** Histograms showing the distribution of substitution outcomes from two DMS studies. The wild-type (WT) Mpro bins are marked by green circles and the bins containing the inactive variant (C145A) are marked by a magenta circle. **(a)** Data from the 2022 growth assay^49^; the error range for replicate WT samples is shown as a green horizontal bar. **(b)** Data from the 2024 FRET assay^55^ show a shifted distribution, with more resolution around the WT value and identification of hyperactive variants. The error range for the WT variant was smaller than the bin width. **(c, d)** Distributions of substitution outcomes reported by each study. **(e, f)** Position classifications from each dataset were assigned using the RheoScale calculator^56^. In the dataset from the 2022 growth assay **(e)**, positions were categorized into five substitution sensitivity classes: Neutral, Moderate rheostat, Rheostat, Adverse, and Toggle, as defined in **Supplementary table 1**. In the dataset from the 2024 FRET assay^55^ **(f)**, more positions were classified as Rheostat or Toggle, fewer positions were classified as Neutral, and four positions fall into a sixth class, “Enhancing” (defined in **Supplementary table 1**). **(g, h)** The spatial distributions of substitution sensitivity classes from the 2022 growth assay **(g)** and the 2024 FRET assay^55^ **(h)** are shown using a model of the Mpro surface (PDB: 5R7Y^64^). The color scheme follows the legend in panels **(e)** and **(f)**. The positions in tan had <5 substitution outcomes reported and thus a meaningful assignment could not be made. Figures were rendered using UCSF Chimera^90^. **Supplementary figure 3** shows additional ball-and-stick representations of these classes so that the buried positions can be visualized.

A similar strategy was used in 2024 for a variation of the FRET-based assay.^55^ Compared to the 2022 FRET study, the 2024 assay^55^ used a shorter induction time for Mpro expression, which would diminish the amount of Mpro protein present in the cell. As a result, the range of measurable values also shifted: The weakly active variants were better discriminated in the 2022 study; the WT-like and hyperactive variants were better discriminated in the 2024 study (**Supplementary figure 2**). The differences observed between these two datasets support our hypothesis that altered *in vivo* abundance can alter DMS results.

Here, we further tested whether single Mpro substitutions can also alter *in vivo* abundance, and whether individual Mpro positions separately affect function (proteolytic activity) and abundance. To identify candidate positions for our work, we used data from the DMS growth assay^49^ to classify each Mpro position based on its overall substitution sensitivity (**Figure 1, Supplementary figure 3, Supplementary table 1**). To assign this behavior, we used the RheoScale calculator^56^: For each protein position, this analytical tool uses the fractions of substitutions that exhibit WT-like, inactive, or intermediate outcomes to determine the position’s overall substitution sensitivity. Examples for six classes of substitution sensitivities are shown in **Supplementary figure 4**. This classification scheme allowed us to identify the set of rheostat positions that is not readily apparent from the average values of a position substitution outcomes.^8^

Of the rheostat positions identified in the 2022 dataset, we selected four positions (48, 52, 77, and 234) for which previously-published computations predicted high/moderate flexibility^57^ **(Supplementary figure 5)**. We also chose two neutral positions from the 2022 growth assay^49^ (50 and 110) with different flexibilities, to serve as a comparison set for positions with fewer intermediate outcomes. At each of these six positions, we made ten random amino acid substitutions, which we previously showed was sufficient to assess a position’s substitution sensitivity class.^7^

### Mammalian cell-based assays of MPro function

To dissect the effects of the 60 single amino acid substitutions on Mpro function and *in vivo* abundance, we designed HEK293 cell-based assays **(Supplementary figure 6)**. A plasmid expressing a cyclic-luciferase reporter enzyme, engineered to contain an Mpro cleavage site, was stably integrated into HEK293 cells to enable constitutive reporter expression. The cyclic-luciferase enzyme expressed from this gene was inactive unless cleaved by Mpro. Cells were then transiently transfected with Mpro substitution variants and the luciferase assay was performed with cell lysate (**Supplementary figure 7**, first column). Since the values measured in the lysate could be sensitive to changes in either Mpro’s proteolytic activity or its cellular abundance, these luciferase assay outputs are equivalent to DMS growth outcomes^49^. In our assays, the “combined effects” for the set of 60 substitutions sampled >90% of the accessible range of luciferase outputs **(Figure 2a),** as defined by the inactive C145A substitution and the maximum observed outcome (V77S, **Figure 3a**).

**Figure 2.**
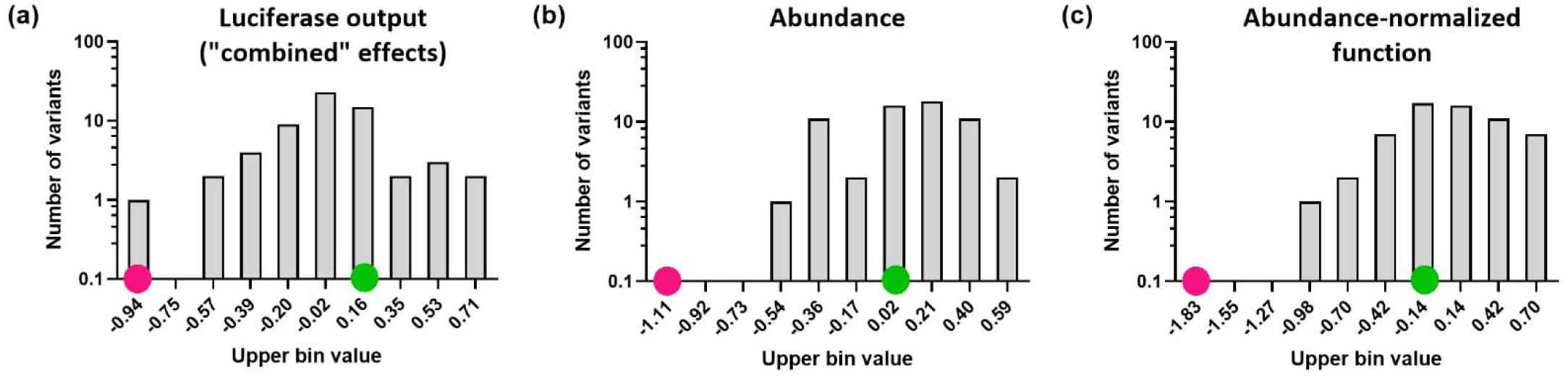
Outcomes measured for the 60 single amino acid Mpro substitutions assessed in this study. **(a)** Luciferase assay output, which measured the combined effects on function and *in vivo* abundance; **(b)** abundance changes were measured by indirect sandwich ELISA; **(c)** function changes were computed by normalizing the measured combined effects with abundance changes. The lower bounds for these three ranges are delineated by inactive Mpro (C145A; a and c) or no protein (b) and are shown with magenta circles. The interval encompassing replicate assays for WT-like variants is marked by the green circle. Note that all three assays showed substitutions with gain-of-function outcomes.

**Figure 3.**
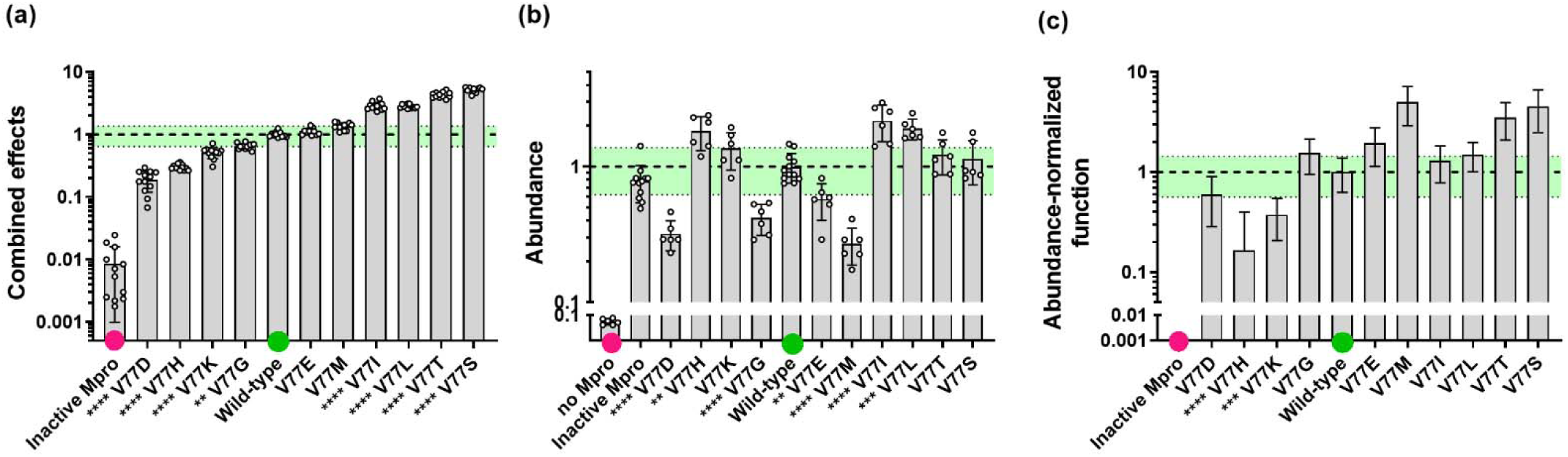
Outcomes measured for the set of substitutions at Mpro position 77. **(a)** Combined effects of substitutions were assessed using a luciferase reporter assay. **(b)** Abundance changes were independently measured using indirect sandwich ELISA. **(c)** Measurements for combined effects **(a)** and abundance changes **(b)** were used to compute the abundance-normalized function for each substitution. Note that the X-axes for the three plots are ordered the same. For all three data sets, substitution outcomes were normalized to the WT value (green circles; dashed line at y=1). The light green regions delineate the standard deviation of all replicate WT values measured during this study. Values corresponding to the inactive variant/no protein are indicated with magenta circles. For each substitution, six technical replicates for each of two biological replicates were measured (open circles); bars show the average for each variant and error bars are standard deviations. Error propagation for (c) is described in Methods. Variants with outcomes statistically different from WT were identified using one-way ANOVAs with Dunnett’s correction (****p <0.0001, ***p<0.001, **p<0.01, *p<0.1); significance asterisks are shown on the X-axis labels. Data for the other five Mpro positions are shown in **Supplementary figure 7** and average values for all variants are available in **Supplementary data**.

When the data from the two DMS studies ^49; 55^ and the current study were compared, the values for the 60 substitution variants showed reasonable correlations **(Supplementary figure 8)**. Differences between the two DMS and our experiments could arise from one or more sources, including: (i) The DMS assays were performed in yeast, whereas our assays were performed in human HEK293 cells. The sensitivity and homeostatic response of these two different cell types to the cleavage of essential cellular proteins is likely to be quite different and could include regulating Mpro expression (ii) The Mpro cleavage site is present in many endogenous proteins, which will differ in the two cell types and could lead to different cytotoxicities. (iii) Indeed, the 2022 yeast growth assay leveraged the toxicity of overexpressed Mpro to measure the effects of substitutions^49^. In contrast, our experiment optimized the concentration of Mpro transfection to allow measurements for a wide range of catalytic activities **(Supplementary figure 9)**. Nevertheless, any variants that increase abundance >4-fold in our experiments might also cause cytotoxicity, which would manifest as diminished activity (**Supplementary figure 9**). (iv) Mpro can cleave a variety of substrate sequences^58^, and the set of endogenous yeast proteins cleaved in the 2022 growth assay^49^ likely includes a variety of substrate sequences. Thus, Mpro variants that altered substrate specificity could have non-linear outcomes on growth. In contrast, our study and the 2024 FRET assay^55^ tested effects on only one substrate sequence.

Next, we compared each *position*’s overall substitution sensitivity among the three studies. Our substitution sensitivities matched those observed in the DMS growth assay^49^ for both neutral positions (50 and 110) and all four rheostat positions (48, 52, 77 and 234) (**Table 1; Supplementary figure 7**, first column). In the 2024 FRET assay^55^, positions 50 and 110 were instead classified as rheostat and moderate rheostat positions, respectively.

**Table 1.** Substitution sensitivities for the various measured outcomes.^i^.

| Position # | 2022 Growth assay<sup>ii</sup> |  |  |  | 2024 FRET assay<sup>iii</sup> |  |  |  | Combined effects from luciferase output (this study) |  |  |  | Abundance (this study) |  |  |  | Abundance-normalized function (this study) |  |  |  |
| --- | --- | --- | --- | --- | --- | --- | --- | --- | --- | --- | --- | --- | --- | --- | --- | --- | --- | --- | --- | --- |
|  | R<sup>iv</sup> | N<sup>v</sup> | T<sup>vi</sup> | C<sup>vii</sup> | R | N | T | C | R | N | T | C | R | N | T | C | R | N | T | C |
| 48 | <u>0.50</u> | 0.40 | 0.00 | R | <u>0.74</u> | 0.16 | 0.00 | R | <u>0.50</u> | 0.66 | 0.00 | R | <u>0.61</u> | 0.40 | 0.00 | R | <u>0.50</u> | 0.30 | 0.00 | R |
| 50 | 0.04 | <u>1.00</u> | 0.00 | N | 0.13 | 0.26 | 0.00 | M | 0.13 | <u>1.00</u> | 0.00 | N | 0.35 | 0.62 | 0.00 | M | 0.39 | <u>0.74</u> | 0.00 | N |
| 52 | <u>0.63</u> | 0.00 | 0.00 | R | <u>0.57</u> | 0.00 | 0.26 | R | <u>0.56</u> | 0.50 | 0.00 | R | 0.39 | 0.60 | 0.00 | M | 0.43 | 0.10 | 0.00 | M |
| 77 | <u>0.53</u> | 0.30 | 0.00 | R | <u>0.65</u> | 0.05 | 0.32 | R | <u>0.78</u> | 0.20 | 0.00 | R | <u>0.61</u> | 0.40 | 0.00 | R | <u>0.61</u> | 0.40 | 0.00 | R |
| 110 | 0.33 | <u>0.75</u> | 0.10 | N | 0.44 | 0.47 | 0.16 | M | 0.30 | <u>0.70</u> | 0.10 | N | 0.22 | <u>1.00</u> | 0.00 | N | 0.26 | <u>0.70</u> | 0.00 | N |
| 234 | <u>0.53</u> | 0.50 | 0.00 | R | 0.39 | 0.53 | 0.11 | M | <u>0.74</u> | 0.30 | 0.00 | R | 0.26 | <u>0.70</u> | 0.00 | N | 0.35 | 0.50 | 0.00 | M |

<sup>i</sup> The averages and standard deviations for each substitution, which were used to determine each position’s sensitivity class, are available in the **Supplementary data**.
<sup>ii</sup> Growth assay results from Flynn *et al.*, (2022) were analyzed using RheoScale (Hodges *et al.*, 2018).
<sup>iii</sup> FRET assay results from Flynn *et al.*, (2024) were analyzed using RheoScale (Hodges *et al.*, 2018).
<sup>iv</sup> Columns labeled “R” contain the RheoScale rheostat scores; positions with RheoScale rheostat scores ≥0.5 (underlined) are defined as “rheostat” positions (Hodges *et al.*, 2018).
<sup>v</sup> Columns labeled “N” contain the RheoScale neutral scores; positions with RheoScale neutral scores ≥0.7 (underlined) are defined as “neutral” positions (Martin *et al.*, 2020).
<sup>vi</sup> Columns labeled “T” contain the RheoScale toggle scores; positions with RheoScale toggle scores ≥0.64 (underlined) are defined as “toggle” positions (Miller *et al.*, 2017).
<sup>vii</sup> The columns labeled “C” contain the substitution sensitivity class for each position. These class assignments are based on the framework outlined in **Supplementary table 1**; letters correspond to: R, rheostat; N, neutral; M, moderate rheostat. No toggle, adverse or enhancing positions were included in the current study.

### The relationship between effects on abundance and function is complex

To determine the contributions of altered *in vivo* protein abundance to changes in luciferase outputs, we independently measured Mpro abundance using an indirect sandwich enzyme-linked immunosorbent assay (ELISA) **(Supplementary figure 6 and Supplementary figure 7)**. Changes in *in vivo* protein abundance can arise from effects on translation rates, folding efficiency, stability, solubility, and degradation pathways. Furthermore, the ELISA does not differentiate between functional homodimers and non-functional monomers. Dimerization is required for Mpro’s catalytic activity^59^ and substitutions that alter the dimer←→ monomer equilibrium would alter the measured function. In addition, substitutions that alter dimerization would also diminish stability, by shifting the folding equilibrium (dimer ←→ monomer ←→ unfolded) to the right, resulting in more unfolded Mpro. Although the unfolded protein could be detected by the ELISA antibody, which targets a linear Mpro epitope (per the manufacturer’s information), we expect that unfolded protein would be quickly degraded (or aggregate) and thus diminish its *in vivo* abundance.

We used Mpro abundance values to normalize the luciferase output values and thereby isolate substitution effects on Mpro’s protease activity. Results from the abundance normalized-function might reflect changes in enzyme catalytic rates, substrate binding affinity, ligand specificity, and/or dimerization. The three outcomes measured in this study are shown in **Figure 2** for the set of 60 substitutions, and example plots for position 77 are shown in **Figure 3**. Data for all other positions are shown in **Supplementary figure 7**.

Even though the un-normalized luciferase output suggested that several substitutions had very low biological activity **(Figure 2a),** no substitutions showed extremely low *in vivo* abundance **(Figure 2b)** or was catalytically inactive **(Figure 2c)**. The set of 60 substitutions showed an ∼45-fold range of changes in abundance **(Figure 2b)** and includes substitutions that produce both lower and higher protein levels than WT. The abundance normalized-function sampled only 60% of the accessible functional range and included substitutions with enhanced function **(Figure 2c).**

Notably, for any of the six positions tested, the rank-order of substitution outcomes for the un-normalized luciferase output does *not* match the order for abundance or normalized-function **(Figure 3; Supplementary figure 7)**. For example, at position 77, substitutions D, G, and M showed a significant decrease in abundance, whereas the normalized-functions of these substitutions were within the range of WT values **(Figure 3)**. Conversely, at position 234, substitutions V and K exhibited significantly lower functional outcomes compared to the WT, although their abundances were near WT levels **(Supplementary figure 7p-r)**. Indeed, when changes in abundance were compared to changes in the un-normalized luciferase outputs for the full dataset, no correlation was observed **(Figure 4a)**. Thus, for Mpro, changes in protein abundance can significantly modulate the observed biological outcomes (**Supplementary figure 10)**, and in some cases, may even overshadow changes in catalytic activity of the substitution itself.

**Figure 4.**
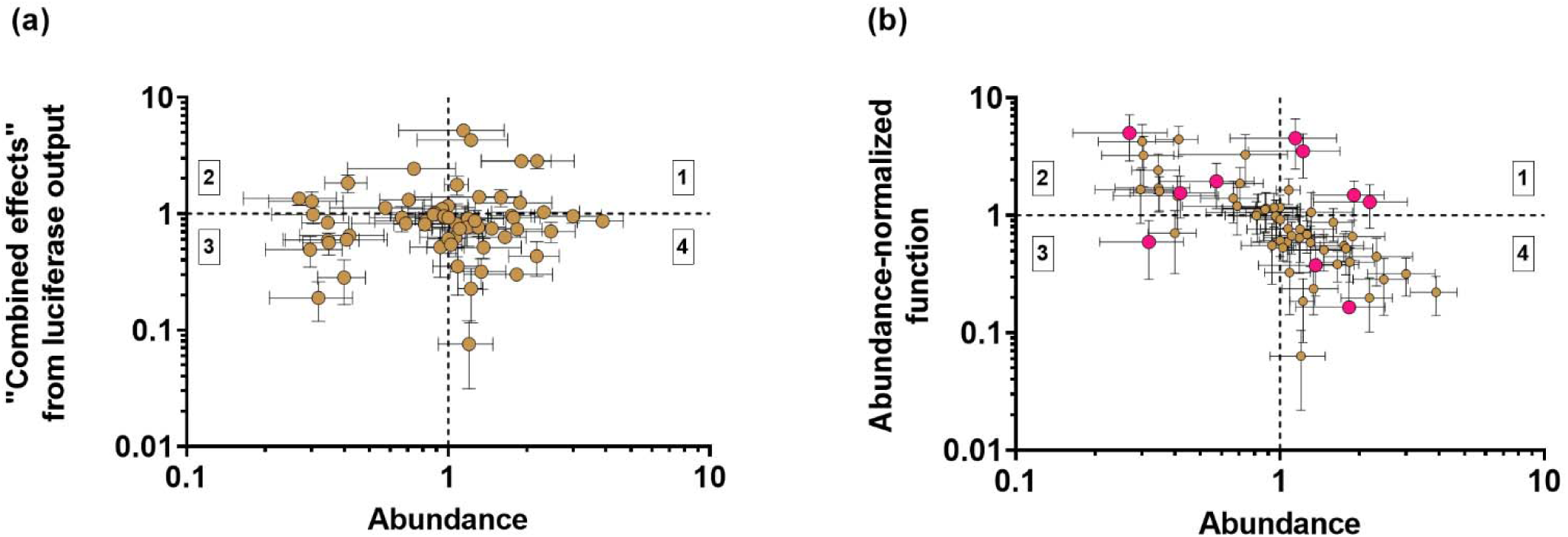
Changes in abundance can make significant contributions to changes in the luciferase assay output. Each data point corresponds to the average values shown in **Supplementary figure 7** and listed in **Supplementary data**; error bars correspond to the standard deviations for each measurement. Dotted lines correspond to the WT values and differentiate the quadrants containing substitutions that are (1) more abundant, more active; (2) less abundant, more active; (3) less abundant, less active; and (4) “more abundant, less active”. **(a)** “Combined effects” of the luciferase assay output versus *in vivo* abundance. The Spearman correlation coefficient is 0.04. **(b)** Normalized function versus *in vivo* abundance. For the whole data set, Spearman r is -0.68 and Pearson r is 0.71, which indicates that enhanced proteolytic activity is often accompanied by diminished Mpro abundance. However, individual plots (**Supplementary figure 11**) show distinct behaviors for individual positions: Positions 48, 50 and 52 show strong relationships, whereas positions 110 and 234 show moderate relationships. Position 77, shown here with large magenta circles, has the weakest correlation (Spearman r = -0.5, Pearson r = 0.39).

Interestingly, a comparison of the changes in abundance and normalized-function revealed a moderate correlation for the set of 60 substitutions (**Figure 4b**). When each of the six positions was examined individually, a strong relationship was observed for positions 48, 50 and 52 (**Supplementary figure 11**). However, this trend was not observed for Mpro position 77 (**Supplementary figure 11, Figure 4b** magenta circles). At this position, individual substitutions showed a variety of outcomes: V77H increased abundance and decreased function, V77M decreased abundance and increased function, V77D decreased both parameters, and other substitutions affected only one of the two parameters.

### Comparisons of substitution outcomes and side chain physicochemical properties

Next, we considered whether the observed changes could be accounted for by the physicochemical properties of the substituted side chains at each position. For each position, we assessed the relationship between its set of substitution outcomes and each of three biophysical traits: secondary structure propensity^60; 61^, side chain size^62^, and hydrophobicity^63^. Several trends emerged.

*Secondary structure propensity* (**Supplementary table 2**). For each Mpro position located in or near an α-helix, we selected the helical propensity score corresponding to its specific or potential location in the helix (*e.g.*, N2, C-cap) as previously defined^59^. For the position located in a β-sheet (position 77), β-sheet propensities were used.^60^ Of the six positions analyzed, positions 52 and 77 showed weak correlations **(Supplementary Figure 12)**. Position 234 exhibited a stronger relationship between helical propensity and normalized function; this suggests that amino acids with higher C2 helical propensity may reduce function. Structural mapping shows that position 234 lies on a surface-exposed helix, distant from the catalytic dyad **(Figure 5)**. This location suggests that position 234 may contribute to functionally important dynamic fluctuations; rigidifying the helix via high-propensity residues could interfere with necessary flexibility.

**Figure 5.**
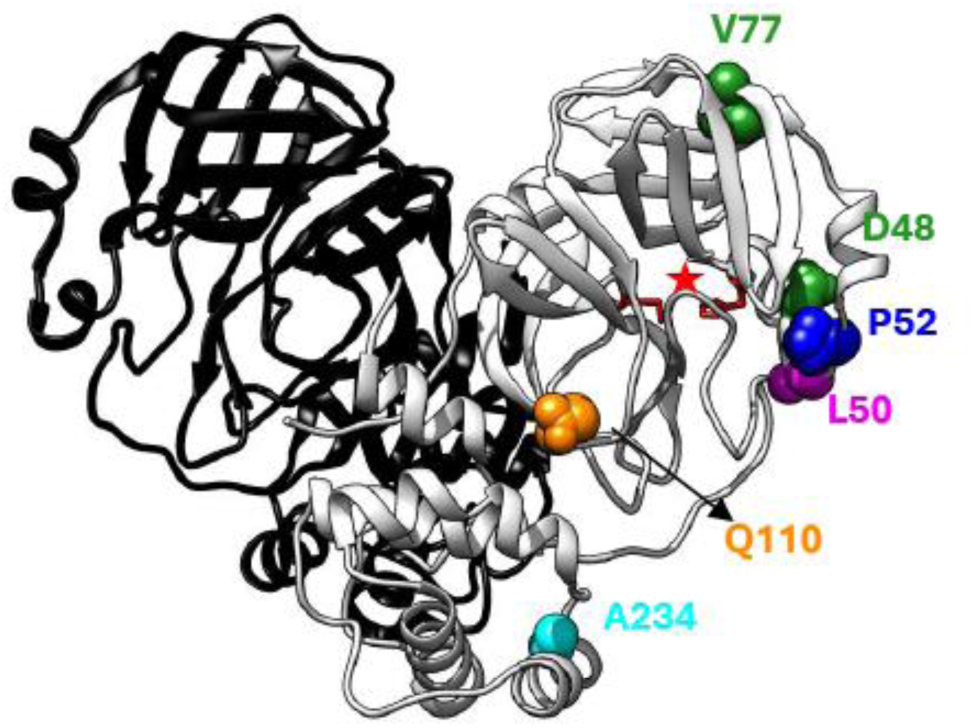
Distinct combinations of substitution sensitivities were observed at the six tested Mpro positions. The six Mpro positions tested in this work are indicated with colored spheres on one monomer of PDB 5R7Y^64^; colors correspond to the different combinations of substitution sensitivities measured for abundance and normalized function **(Table 1)**. The histidine and cysteine side chains forming the catalytic dyad (at positions 41 and 145) are in red and the active site is indicated by the red star. Orange position 110: neutral for both abundance and normalized-function. Green positions 48 and 77: rheostatic outcomes for both abundance and normalized function. Blue position 52: moderate rheostat for both abundance and normalized-function. Magenta position 50: moderate rheostat for abundance and rheostat for normalized-function. Cyan position 234: neutral for abundance and moderate rheostat for normalized-function. Figure rendered using UCSF Chimera.^90^

*Side chain size* (**Supplementary table 2**). We next compared the substitution outcomes with amino acid sizes^62^ **(Supplementary figure 13)**. Two positions – 48 and 77 – exhibited modest trends. At position 48, a positive correlation was observed between side chain volume and normalized function when one outlier (arginine) was excluded. Position 48 is located on the solvent-exposed surface of a helix. Position 48 is near the loop containing the catalytic H41 but is >13 Å from the active site ^64^. Thus, we speculate that substitutions at position 48 alter the dynamics of the active site.

At position 77, when two outliers were excluded (glycine and aspartate), side chain size negatively correlated with the combined effects on the luciferase output and with normalized-function. Position 77 resides on a β-strand, and smaller residues allow enhanced activity. Position 77 is >17 Å from the active site, so we again speculate that substitutions alter dynamics.

*Hydrophobicity.* Finally, we compared substitution outcomes for each position with Wimley–White interfacial hydrophobicity scores^63^ **(Supplementary figure 14)**. At position 48, substitutions with higher hydrophobicity were associated with increased *in vivo* abundance but reduced normalized function. This suggests that changes in hydrophobicity is the source of the trade-off observed between abundance and function (**Supplementary figure 11**). Since position 48 is solvent exposed in the wild-type structure (**Figure 5**), it is easy to rationalize that hydrophobic substitutions are detrimental to abundance. However, the origins of enhanced function remain unclear.

Position 234 displayed the strongest hydrophobicity-associated trends. Hydrophobic substitutions strongly correlated with both higher luciferase output and increased normalized function. This trend may be related to the fact that, in the wild-type structure, the side chain of position 234 makes direct hydrophobic contacts with the side chains of F230 and Y239, and additional hydrophobic residues are in the vicinity.^59; 65^

### Substitution sensitivities of individual Mpro positions reveal distinct combinations of neutral and rheostat behaviors for function and abundance

Finally, we determined the overall substitution sensitivity classes (**Supplementary table 1**) for normalized-function and abundance for the six Mpro positions **(Table 1**; **Figure 5**). Despite examining just six positions, the data reveals a wide range of behaviors. Positions 48 and 77 showed strong rheostat effects for both parameters, whereas position 52 displayed moderate rheostat behavior for both, and position 110 displayed neutral behavior for both. Position 234 was neutral for abundance but rheostat for function, and position 50 was a moderate rheostat for abundance and neutral for function.

The different sensitivities of these six positions are not readily explained by their locations on the Mpro structure **(Figure 5)**. Positions with different types of outcomes are adjacent to each other, and none are near the active site or the dimerization interface. We also considered solvent exposure, secondary structure, and proximity to the catalytic dyad. However, none of these clearly accounted for the observed differences in substitution sensitivities **(Supplementary table 2).** This matches our studies of rheostat positions in other proteins, which also are distributed widely across protein structures and lack obvious relationships to structural features.^1; 7; 40; 66^

## DISCUSSION

In this set of 60 Mpro substitutions, some altered function, some altered abundance – to different magnitudes, and in same or different directions. For the first time, we also identified positions where substitutions simultaneously modulate *both* function and abundance. These two parameters did not readily correlate, hinting that they could be controlled by different biophysical features. Furthermore, the fact that multiple combinations of substitution sensitivities were observed for just six positions suggests that such variability is likely widespread across Mpro positions.

Our result is in striking contrast to the assumption underlying many DMS studies, that single amino acid substitutions do not greatly alter abundance,^11; 67; 68^ but it is in agreement with other studies that demonstrate single substitutions alter abundance in other proteins (e.g., ^18; 45; 47; 69; 70^). For some applications, the distinction between changes in function and abundance might not matter. As noted above, Flynn *et al.*, strove to identify Mpro positions that did not tolerate substitutions. In another example, the DMS study of p53 was used to identify missense mutations that could lead to cancerous phenotypes.^71^ In both of these studies, the “combined effects” from DMS are useful to understand biological outcomes and the mechanism by which the protein was inactivated was irrelevant.

However, to understand the general principles by which proteins *work*, which is critical for developing accurate prediction algorithms, the DMS assay design is not sufficient. Indeed, although the substitution classes for each Mpro position (rheostat or neutral, **Table 1**) for combined effects from DMS/luciferase outputs appear to agree with those for abundance-normalized function, the rank order of substitution outcomes differ. This highlights the fact that different *substitutions* can make diverse contributions to individual protein parameters.

Another complex relationship illustrated by the current results is that protein *positions* can play distinct roles in regulating function and abundance. The examples here for Mpro positions 48 and 77 are, to our knowledge, the first documentation that some protein positions can have rheostatic effects on both parameters. This parallels observations in other proteins, which have rheostat positions that alter multiple *functional* parameters. For example, rheostat positions in human liver pyruvate kinase affected up to five different functional parameters (binding affinity to its substrate, two different allosteric effectors, and two allosteric coupling constants).^72^ In a second example, a subset of rheostat positions in *Escherichia coli* LacI affected both transcription repression and response to allosteric ligand.^73^

Studies of single substitutions in other proteins also show complex relationships between function and abundance, with the two parameters altered to different extents and in different directions.^74; 75^ In the current study, the sets of substitutions for three positions show a clear trade-off, with increased function offset by decreased abundance **(Supplementary figure 11)**. It is interesting to think about the possible mechanisms underlying this trade-off. Abundance changes might derive from changes in protein stability, which has been observed in many studies^47; 76; 77^ and is often expected in protein engineering^78–81^. Alternatively, the function/abundance trade-off could arise from a cellular response, with the HEK293 cells limiting the overall level of Mpro activity.

However, we reason that a cell-based response should equally affect all substitutions at all positions. In contrast, the stability/function trade-off is not universal.^34; 45; 82; 83^ Thus, the *lack* of correlation for Mpro position 77 supports the hypothesis that positions 48, 50, and 52 experience a stability/function trade-off. These results lead us to further hypothesize that some positions may act as rheostats for both function and stability. Prior to this work, stability rheostat positions were known for (i) *Zymomonas mobilis* pyruvate kinase, but they had few detectable effects on function^84; 85^ and (ii) GB1^1; 86^, for which effects on function are not known.

In conclusion, this study illustrates that single substitutions can separately affect different protein parameters. Each parameter can have different biophysical bases, which likely requires a unique model to predict. Ultimately, integrating parameter-specific contributions is critical to build biophysical models that accurately capture actual protein behavior.

## MATERIALS AND METHODS

### Identification of rheostat positions in the Mpro DMS dataset

Raw data from the previously published deep mutational scanning (DMS) datasets for the main protease (Mpro) of SARS-CoV2^49; 55^ (from the supplementary file associated with the 2022 Growth assay^49^ and Supporting Information available for the 2024 FRET assay^55^) was analyzed using the RheoScale calculator^56^. For these analyses, the lower limit of measurable substitution outcomes assay was determined from values for the nonsense mutations corresponding to positions 8 through 50, the upper limit was set by the maximum outcome measured, and the bin number was set to 10. The rheostat, toggle, and neutral scores thus obtained were used to assign the position types as discussed in **Supplementary table 1**. Such class assignment for all Mpro positions from the DMS growth assay^49^ is available in **Supplementary figure 1c**.

### DNA constructs

The plasmid pCS-NSP5, which expresses wild-type Mpro (Uniparc ID: UPI00137481FB) from the CMV promoter, was used to generate Mpro variants. For each targeted amino acid position, 10 randomly chosen, single amino acid substitutions were produced by Twist Biosciences (San Francisco, CA). Plasmids were transformed into DH5-alpha competent cells (Catalog # C2987H, New England Biolabs Ipswich, MA), and purified using QIAprep Spin Mini Kit (Catalog #27104, Hilden, Germany).

The cyclic-luciferase reporter plasmid pCycLucNSP5, which was used to assay Mpro activity, was created by modifying the plasmid cycLucTEVS (a gift from Roman Jerala, Addgene plasmid #119207).^87^ This included replacing the cleavage site for the tobacco etch virus protease with the cleavage site for SARS-CoV2 Mpro (SAVLQ↓SGFRK)^58^ and removing the PEST sequence degron from the C-terminus of the cycLUC.

### Cell culture

HEK293 cells (American Type Culture Collection, Manassas, VA; catalog number CRL-1573) were cultured in Dulbecco’s Modified Eagle Medium with 4.5 g/L glucose, supplemented with 10% fetal bovine serum, 100 µg/mL penicillin, and 100 U/mL streptomycin. Stable cell lines expressing the reporter were maintained in 300 µg/ul G418 (CAS # 108321-42-2; Goldbio, St. Louis, MO). Cells were maintained at 37 °C under a humid atmosphere containing 5% CO_2_. Growth was monitored daily, and cells were split at 80-90% confluency. The growth media was changed as required. Cells were used for a maximum of 20 passages. Subcultures for transfection were prepared in 96-well plates with 2.5×10^4^ cells/well and 200 µL complete medium.

### Stable cell line generation

The reporter plasmid pCycLucNSP5 was transfected into HEK293 cells using Turbofect transfection reagent (Thermo Fisher Scientific, Waltham, MA). Selection for stable expression of the plasmid was performed with 600 µg/ml G418. Multiple clones (HEK293-CLucNSP5) were isolated and utilized in experiments.

### Transfection of Mpro substitutions to assay protease activity

HEK293-CLucNSP5 cells were transfected in 96-well plates utilizing Turbofect transfection reagent (Thermo Fisher Scientific, Waltham, MA). Unless otherwise stated, each transfection contained 15 ng of pCS-NSP5 expressing WT or variant Mpro, 10 ng pTK-Renilla (when encoded *Renilla* luciferase that was used as a transfection efficiency control), and 175 ng pCMV-LacZ (as stuffer plasmid). Culture media was changed four hours post transfection. Cells were incubated at 37 °C overnight before performing the luciferase assay or ELISA.

### Luciferase assay

Luciferase was measured using the Bright-Glo™ Luciferase Assay System kit (Catalog # E2620) per manufacturer’s instructions except as noted below, and results were assayed using a Tecan plate reader (Mannedorf, Zurich, Switzerland). A modification of the reagent volumes for the luciferase assay was employed, using 50 µl of LAR II reagent per well for firefly luciferase activity. After measuring firefly luciferase signal, 50 µl of 1X Stop & Glo reagent was added per well to quench the firefly luciferase signal and measure *Renilla* luciferase activity.

To optimize the linear range of luciferase response, we titrated the WT Mpro plasmid used for transfection, from 2.5 ng to 100 ng per well. Luciferase output increased linearly up to 60 ng and plateaued/decreased at higher concentrations **(Supplementary figure 8)**. We conclude that higher doses are associated with cytotoxicity. Based on these results, 15 ng plasmid was selected for all subsequent experiments, to ensure that measurements for Mpro substitution variants were within the assay’s dynamic range.

Raw data for all Mpro substitutions tested in this study have been deposited in https://github.com/ShwethaSreenivasan/Mpro_abundance_vs_function/blob/main/Mpro_variant_data_22Dec25.xlsx.

### ELISA to measure Mpro abundance changes

HEK293-CLucNSP5 cells transfected with Mpro were lysed with 25 µl of 1 X ELISA Cell Extraction Buffer (Catalog #69905, Cell Signaling Technology, Danvers, MA) per well in a 96-well plate. Indirect sandwich ELISA was performed using a kit from Elabscience® (Catalog # E-ELIR-K001, Houston, TX) in clear 96-well ELISA plates (Catalog # 15031, Thermo Fisher Scientific, Waltham, MA). The plates were coated with SARS-CoV-2 3C-Like Protease capture antibody (Catalog #51661a, Cell Signaling Technology, Danvers, MA) at 1 µg/mL in coating buffer and incubated at 4 °C overnight. Wells were washed and blocked, and 10 µl of cell lysate (diluted 1:3 in the blocking buffer) was added. For detection, 2 ng/mL Strep II tag antibody (Catalog # A01737-100, GenScript, Piscataway, NJ) was added to wells and incubated at room temperature for two hours with agitation. Following washing, goat anti-mouse IgG (H + L)-HRP Conjugate (Catalog #1706516, Bio-Rad, Hercules, CA) was added and incubated for an hour at room temperature, followed by addition of 100 µL of TMB substrate to each well for detection. Following incubation for 12-15 minutes, stop solution was added and absorbance at 450 nm measured within 30 minutes using a Tecan Plate Reader (Mannedorf, Zurich, Switzerland). Raw data have been deposited in https://github.com/ShwethaSreenivasan/Mpro_abundance_vs_function/blob/main/Mpro_variant_data_22Dec25.xlsx.

### Computing abundance normalized-function

Three experimental measurements were made for each Mpro WT and variants (i) Mpro’s *in vivo* activity was measured using the firefly luciferase activity. (ii) Transfection differences were controlled by measuring the *Renilla* luciferase signal intensity from the control plasmid (pTK-Renilla). (iii) Mpro abundance was determined from ELISA. For each variant, 4-6 technical replicates for two biological replicates were assayed. These three measurements were then used to generate three different outcomes for each Mpro variant.

*Mpro outcome 1: Combined effects*. For each Mpro variant, the “combined effect” was obtained by normalizing the firefly luciferase activity to the *Renilla* luciferase activity. This was then normalized to the average of the values from the wells containing Mpro WT replicates that were present in each plate, with appropriate error propagation (Equation 1 below), to control day-to-day technical variation.

*Mpro outcome 2: Abundance changes*. The abundance changes for each Mpro variant were obtained by normalizing results from the indirect ELISA assay to each plate’s WT average (same as in Mpro outcome 1), with appropriate error propagation (Equation 1 below), to control for day-to-day technical variation.

*Mpro outcome 3: Abundance normalized-function*. Abundance normalized-function for each variant was obtained by calculating the ratio of the normalized combined effects to normalized abundance changes, with appropriate error propagation (Equation 1).

For all error propagation during normalizations, we used the following formula:

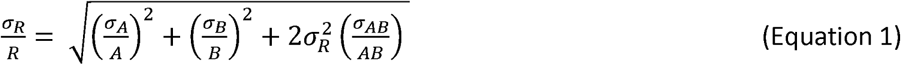

where, *R* is the ratio of outcomes *A* and *B*, and *σ_R_*, *σ_A_* and *σ_B_* are their respective standard deviations; *σ_AB_* is determined by ratio of the standard error of means of *A* and *B* as reported by GraphPad Prism version 9.4.1.

The independent measurements for these three outcomes are reported in https://github.com/ShwethaSreenivasan/Mpro_abundance_vs_function/blob/main/Mpro_variant_data_22Dec25.xlsx. All data plots were made using GraphPad Prism version 9.4.1.

### Statistical analyses of experimental data

To identify which variants differed from WT, statistical analyses were carried out using GraphPad Prism version 9.4.1. The data acquired followed a log-normal distribution, as indicated by D’Agostino-Pearson tests. Thus, all values were transformed to log10 prior to statistical tests. One-way ANOVA was performed with Dunnett’s correction for multiple comparison of each substitution at a position to the value collected for WT of that position’s data set; statistically significant variants were identified based on Dunnett’s correction (****p <0.0001, ***p<0.001, **p<0.01, *p<0.1); complete reports of these statistical analyses are included in **Supplementary table 3**.

### Assigning overall substitution outcomes to each tested Mpro position

The measurements for combined effects, abundance-normalized function, and abundance changes were assessed using the RheoScale calculator to obtain the rheostat, toggle, and neutral scores for each of these parameters at the seven positions tested in this study **(Table 1)**. For these analyses, the histogram ranges were determined by the minimum and maximum values for each of the three outcomes measured in this study, and the bin number was set to 10. The scores used to assign the position types are described in **Supplementary table 1**.

## DATA AVAILABILITY

The number of replicates, averages and standard deviations for the values shown in **Figure 3** and **Supplementary figure 7** are available on https://github.com/ShwethaSreenivasan/Mpro_abundance_vs_function/blob/main/Mpro_variant_data_22Dec25.xlsx

## SUPPLEMENTARY DATA

Supplementary Data are available at Protein Science. Additional citations associated with these data include^78–81; 88; 89^.

## ACKNOWLEDGEMENTS

We thank Halya Fedosyuk (KUMC) for assistance with initial experiments and Dr. Aron Fenton, Dr. Pierce O’Neil, Anastasiia Sivchenko and Carter Grey (KUMC) for discussion on the manuscript. We thank Jordan Baker (KUMC, Department of Biostatistics and Data Science) for providing suggestions for data analyses. Finally, we thank Dr. Paul Campitelli and Dr. Banu Ozkan (Arizona State University) for sharing and discussing their published DFI scores computed for Mpro.

