## Supplementary for "Dissecting the effects of single amino acid substitutions in SARS-CoV2 Mpro"

**Supplementary data:**

Average values and standard errors of the mean for each variant of Mpro shown in **Supplementary figure 7** and **Figure 3** of the main text are posted as an Excel™ file on https://github.com/ShwethaSreenivasan/Mpro_abundance_vs_function/blob/main/Mpro_variant_data_22Dec25.xlsx

**Supplementary figure 1. Substitutions at rheostat positions broadly sample the accessible range of measured values.** The example rheostat position shown is position 2 in SARS-CoV-2 Mpro, as measured in the DMS growth assay by Flynn *et al.,* in[1]. The WT amino acid is glycine and the average value for catalytically-inactive Mpro -0.01 ± 0.04 in this assay. The range of outcomes determined for WT replicates is 0.97±0.16, and the outcome measured for the WT variant at this position is 0.96.

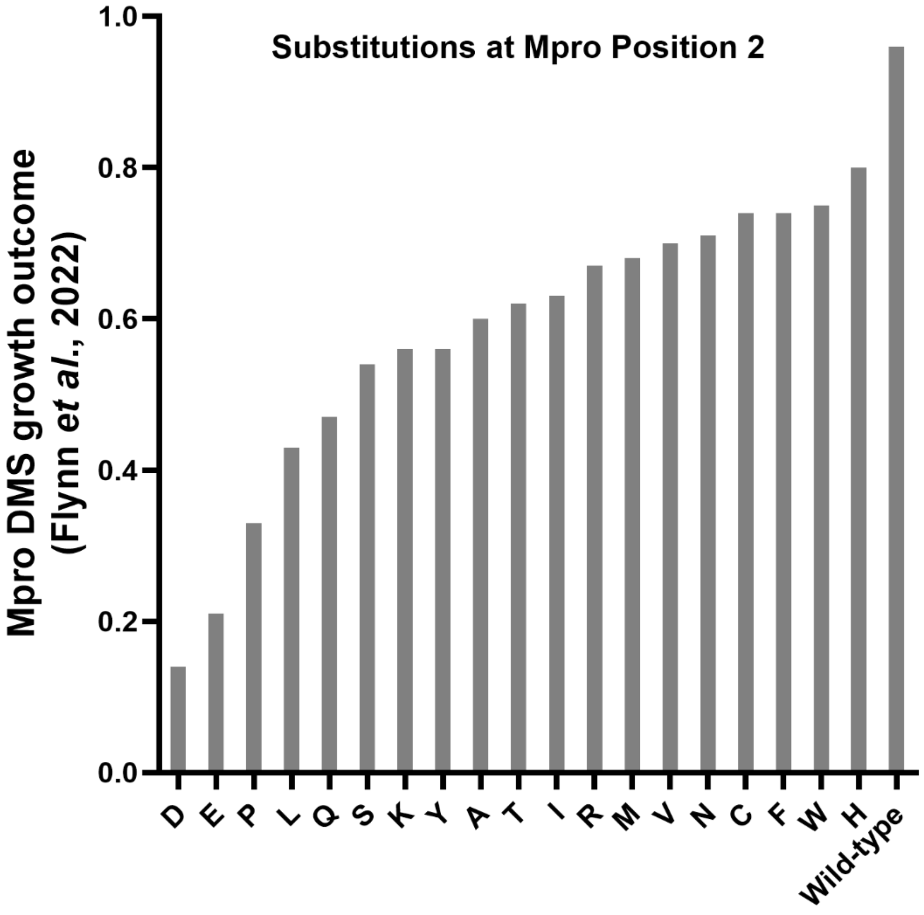

**Supplementary figure 2. Summary of previously published Mpro deep mutational scanning (DMS) data.** For each assay, variants are classified as having: Inactive outcomes – lack of Mpro activity could arise from loss of catalysis and/or from reduced *in vivo* abundance; WT-like outcomes – statistically equivalent to WT (within two standard deviations of the mean of synonymous variants); intermediate outcomes – between inactive and WT-like values; or hyperactive outcomes – more than two standard deviations more active than the WT average. The legends below indicate the number of variants in each of these categories. Note that, conventionally, inactive variants are referred to as "loss-of-function" variants and hyperactive variants are called "gain-of-function" variants. However, since altered biological activity in these assays could result from altered catalysis or *in vivo* abundance (or both), we do not use this nomenclature.

**(a-c)** In their publication from 2022, Flynn *et al.*,[1] reported results for three independent DMS assays of Mpro activity. The distributions of substitution outcomes are shown: **(a)** A yeast growth assay ("growth assay") measured the toxicity of Mpro to yeast growth, which is presumably due to Mpro's cleavage of endogenous yeast proteins. **(b)** A FRET-based assay assessed Mpro activity by measuring the loss of fluorescence transfer between two fluorescent proteins joined with a linker containing the Mpro cleavage site. **(c)** A transcription factor (TF) assay assessed Mpro activity by measuring the proteolytic inactivation of a transcription factor driving GFP expression. In general, results for these assays showed good agreement with each other.[1] Among these three assays, the yeast growth assay **(a)** identified the highest number of variants with intermediate activity and is used as a reference for our current work.

**(d)** In 2024, Flynn *et al.,*[2] repeated the 2022 FRET-based assay with the induction time reduced by half. This design provided better resolution for outcomes near the WT values and hence identified a greater number of intermediate variants and hyperactive variants. Such enhanced outcomes have previously been observed in other proteins, and frequently occur at rheostat positions.[3-7] Importantly, the differences between the 2022 and 2024 FRET results indicate that the assay is sensitive to changes in Mpro abundance.

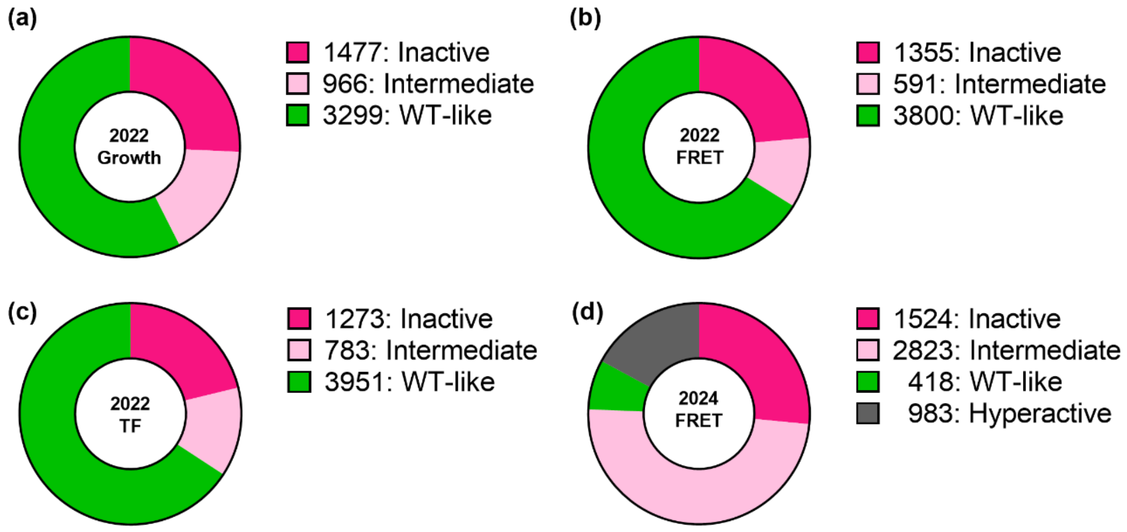


**(e)** Comparison of results from the 2022 growth assay[1] and the 2024 FRET assay[2]. Each data point corresponds to an individual substitution. The shape of this relationship illustrates the effects that arise from their different assay designs. Variants chosen for the current study are shown with orange circles.

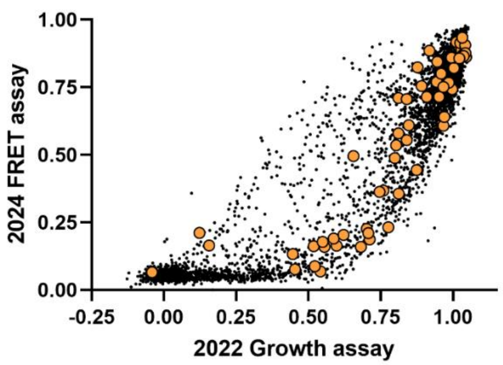

**Supplementary figure 3. Substitution sensitivities classes for Mpro positions derived from the 2022 and 2024 datasets. (a)** As described in the main text Methods, data from the 2022 growth assay and the 2024 FRET assay were used to assign the substitution sensitivity class for each Mpro position:

N – neutral, M – moderate rheostat, R – rheostat, A – adverse, T – toggle, E – enhancing

Two position classes, adverse and enhancing, are defined for the first time in this work. These and other class definitions are further described in **Supplementary table 1**. Differences in position class assignments between the two assays are further discussed in the main text and **Figure 1**.

| | 1 | 2 | 3 | 4 | 5 | 6 | 7 | 8 | 9 | 10 | 11 | 12 | 13 | 14 | 15 | 16 | 17 | 18 | 19 | 20 | 21 | 22 | 23 | 24 | 25 | 26 | 29 | 30 | 31 | 32 | 33 | 34 | 35 | 36 | 37 | 38 | 39 | 40 |
|---|---|---|---|---|---|---|---|---|---|---|---|---|---|---|---|---|---|---|---|---|---|---|---|---|---|---|---|---|---|---|---|---|---|---|---|---|---|---|
| 2022 Growth | N | R | R | R | R | R | T | R | T | T | A | R | R | T | M | R | R | R | M | R | N | R | N | N | R | N | T | R | T | R | N | N | N | R | M | R | T | T |
| 2024 FRET | M | R | T | R | R | T | T | R | T | T | T | T | T | T | R | T | T | R | R | T | R | R | R | M | M | R | T | R | T | T | E | M | N | T | R | R | T | T |

| | 41 | 42 | 43 | 44 | 45 | 46 | 47 | 48 | 49 | 50 | 51 | 52 | 53 | 54 | 55 | 56 | 57 | 58 | 59 | 60 | 61 | 62 | 63 | 64 | 65 | 66 | 67 | 68 | 69 | 70 | 71 | 72 | 73 | 74 | 75 | 76 | 77 | 78 |
|---|---|---|---|---|---|---|---|---|---|---|---|---|---|---|---|---|---|---|---|---|---|---|---|---|---|---|---|---|---|---|---|---|---|---|---|---|---|---|
| 2022 Growth | T | R | R | R | N | N | N | R | R | N | M | R | N | R | N | N | R | M | N | N | M | N | M | N | N | R | N | R | N | N | N | N | N | N | R | N | R | N |
| 2024 FRET | T | A | R | R | M | M | M | R | R | M | M | R | M | T | M | N | R | R | N | M | R | N | M | N | M | R | M | R | N | N | M | N | M | N | R | M | R | M |

| | 79 | 80 | 81 | 82 | 83 | 84 | 85 | 86 | 87 | 88 | 89 | 90 | 91 | 92 | 93 | 94 | 95 | 96 | 97 | 98 | 99 | 100 | 101 | 102 | 103 | 104 | 105 | 106 | 107 | 108 | 109 | 110 | 111 | 112 | 113 | 114 | 115 | 116 |
|---|---|---|---|---|---|---|---|---|---|---|---|---|---|---|---|---|---|---|---|---|---|---|---|---|---|---|---|---|---|---|---|---|---|---|---|---|---|---|
| 2022 Growth | N | M | N | M | N | N | R | N | R | N | R | N | R | N | N | N | R | N | M | R | R | N | M | N | N | N | M | R | N | R | R | N | T | R | T | R | R | R |
| 2024 FRET | M | M | N | R | N | M | R | R | R | N | R | N | A | N | N | N | T | N | M | R | R | M | R | R | R | M | R | R | M | R | T | R | T | R | T | R | T | R |

| | 117 | 118 | 119 | 120 | 121 | 122 | 123 | 124 | 125 | 126 | 127 | 128 | 129 | 130 | 131 | 132 | 133 | 134 | 135 | 136 | 137 | 138 | 139 | 140 | 141 | 142 | 143 | 144 | 145 | 146 | 147 | 148 | 149 | 150 | 151 | 152 | 153 | 154 |
|---|---|---|---|---|---|---|---|---|---|---|---|---|---|---|---|---|---|---|---|---|---|---|---|---|---|---|---|---|---|---|---|---|---|---|---|---|---|---|
| 2022 Growth | R | R | N | R | N | R | M | R | R | R | R | R | R | R | T | R | R | N | R | R | R | R | R | R | N | M | T | R | T | T | T | R | T | T | N | R | N | N |
| 2024 FRET | T | T | R | T | M | T | R | T | T | A | R | R | R | R | T | R | T | M | T | R | R | T | R | T | E | M | T | T | T | T | T | A | T | T | E | R | M | N |

| | 155 | 156 | 157 | 158 | 159 | 160 | 161 | 162 | 163 | 164 | 165 | 166 | 167 | 168 | 169 | 170 | 171 | 172 | 173 | 174 | 175 | 176 | 177 | 178 | 179 | 180 | 181 | 182 | 183 | 184 | 185 | 186 | 187 | 188 | 189 | 190 | 191 | 192 |
|---|---|---|---|---|---|---|---|---|---|---|---|---|---|---|---|---|---|---|---|---|---|---|---|---|---|---|---|---|---|---|---|---|---|---|---|---|---|---|
| 2022 Growth | M | N | R | N | T | M | R | R | T | R | R | R | R | N | M | R | M | R | R | T | R | R | R | N | T | N | R | R | T | M | R | R | T | R | R | R | N | R |
| 2024 FRET | R | M | A | M | T | R | T | A | T | A | T | R | T | M | R | R | R | T | T | T | T | T | R | N | T | N | T | T | T | R | T | R | T | R | R | R | M | T |

| | 193 | 194 | 195 | 196 | 197 | 198 | 199 | 200 | 201 | 202 | 203 | 204 | 205 | 206 | 207 | 208 | 209 | 210 | 211 | 212 | 213 | 214 | 215 | 216 | 217 | 218 | 219 | 220 | 221 | 222 | 223 | 224 | 225 | 226 | 227 | 228 | 229 | 230 |
|---|---|---|---|---|---|---|---|---|---|---|---|---|---|---|---|---|---|---|---|---|---|---|---|---|---|---|---|---|---|---|---|---|---|---|---|---|---|---|
| 2022 Growth | N | R | R | N | T | N | N | R | R | M | T | R | R | R | R | R | R | R | R | M | R | M | M | R | N | M | R | N | N | N | N | N | N | M | N | N | N | R |
| 2024 FRET | M | R | R | M | T | R | M | T | R | R | T | R | A | T | R | R | R | T | R | M | R | R | M | R | N | M | R | M | N | N | N | N | M | M | M | N | M | R |

| | 231 | 232 | 233 | 234 | 235 | 236 | 237 | 238 | 239 | 240 | 240 | 241 | 242 | 243 | 244 | 245 | 246 | 247 | 248 | 249 | 250 | 251 | 252 | 253 | 254 | 255 | 256 | 257 | 259 | 260 | 261 | 262 | 263 | 264 | 265 | 266 | 267 | 268 |
|---|---|---|---|---|---|---|---|---|---|---|---|---|---|---|---|---|---|---|---|---|---|---|---|---|---|---|---|---|---|---|---|---|---|---|---|---|---|---|
| 2022 Growth | R | N | N | R | N | N | N | N | N | N | R | N | A | N | N | N | N | M | N | N | R | N | N | R | N | N | N | R | R | N | R | N | N | R | R | N | M | R |
| 2024 FRET | R | N | N | M | N | N | N | M | M | N | R | M | A | M | N | N | M | M | M | N | R | M | E | R | M | M | M | R | R | N | R | M | N | R | M | N | M | A |

| | 269 | 270 | 271 | 272 | 273 | 274 | 275 | 276 | 277 | 278 | 279 | 280 | 281 | 282 | 283 | 284 | 285 | 286 | 287 | 288 | 289 | 290 | 291 | 292 | 293 | 294 | 295 | 296 | 297 | 298 | 299 | 300 | 301 | 302 | 303 | 304 | 305 | 306 |
|---|---|---|---|---|---|---|---|---|---|---|---|---|---|---|---|---|---|---|---|---|---|---|---|---|---|---|---|---|---|---|---|---|---|---|---|---|---|---|
| 2022 Growth | N | N | R | R | N | N | R | N | N | M | N | N | R | R | M | N | M | M | N | T | A | T | R | R | N | N | T | R | N | R | R | R | N | M | N | M | M | N |
| 2024 FRET | N | N | R | M | N | N | R | M | N | M | M | M | R | R | R | M | R | R | M | T | T | T | R | R | M | M | T | T | M | T | T | R | M | M | M | M | M | N |

**Supplementary figure 3b (next page).** Structural representation of position class assignments on the Mpro structure (PDB: 5R7Y[8]). For each dataset, positions are colored by their position classes using the same color scheme as in panel **(a)**. The left column shows assignments based on the 2022 growth assay; the right column shows those from the 2024 FRET assay. The top row displays neutral and enhancing positions (dark green ball-and-stick and blue space-filling, respectively), the middle row displays moderate rheostat and rheostat positions (ball-and-stick in light green and pink, respectively), and the bottom row shows adverse and toggle positions (light space-filling and dark magenta ball-and-stick, respectively). These visualizations reveal that all position classes are broadly distributed on the protein structure.

**Supplementary figure 3b.**

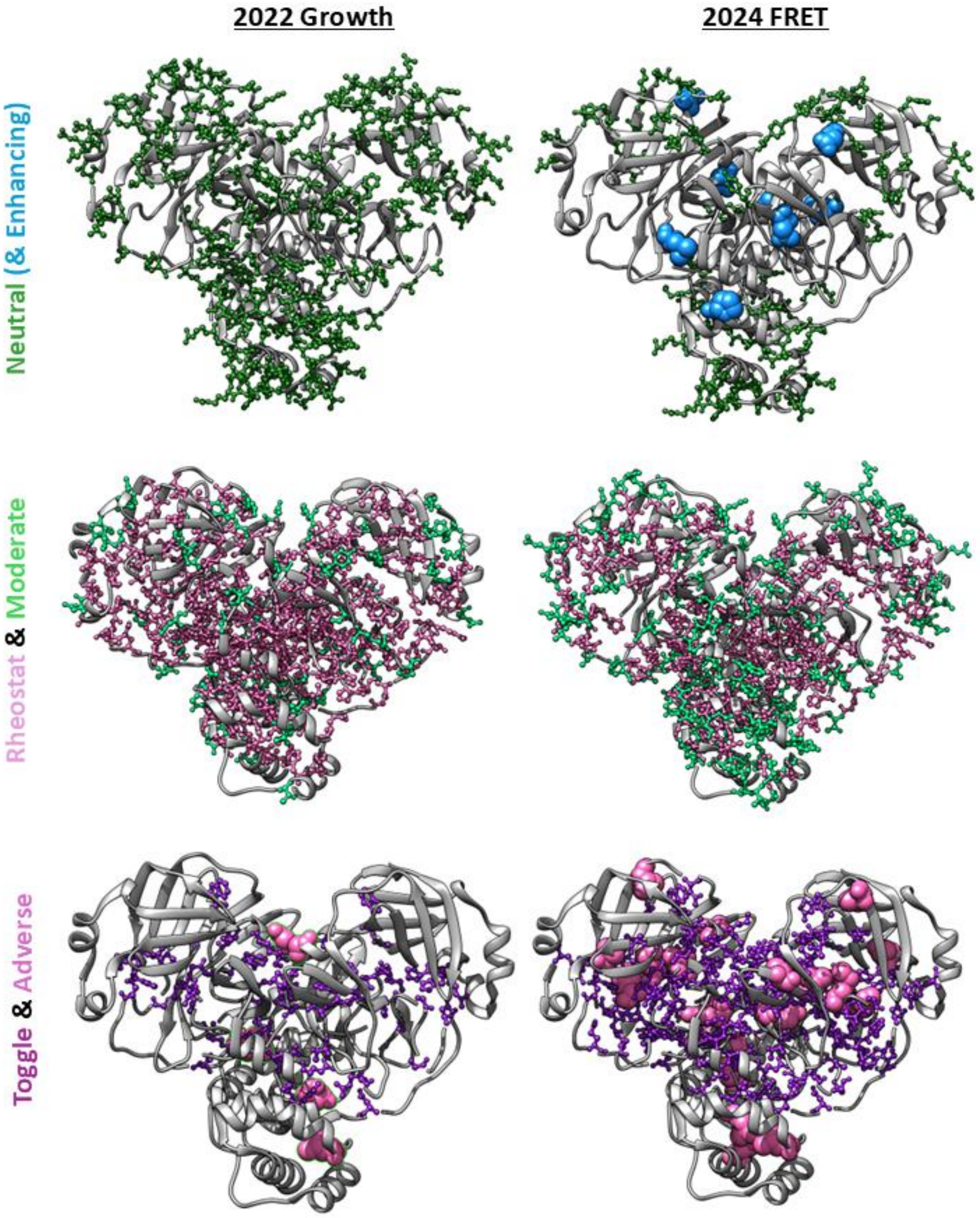

**Supplementary table 1. Framework to classify Mpro positions based on their substitution sensitivities.** Each position's overall sensitivity was classified based on results for multiple amino acid substitutions at that position. (DMS studies substitute all 20 possible amino acids at each position.) Calculations use a histogram-based analysis (RheoScale[9]) to generate three scores – neutral, rheostat, toggle – that quantify how the outcomes sample the accessible range of the measured parameter. Position classes are then assigned using empirically-determined thresholds; these are extensively discussed in the citations listed in the table.

| Position class | Definition | RheoScale scores | Number of Mpro positions | |
|---|---|---|---|---|
| | | | 2022 Growth | 2024 FRET |
| ENHANCING | >80% of the substitutions at a position exhibit activity greater WT. This category was defined during this work. | Enhancing score >0.8 | 0 | 4 |
| NEUTRAL | >70% of substitutions at position behave like WT [10] | Neutral score > 0.7 | 113 | 43 |
| MODERATE RHEOSTAT | Substitution outcomes are statistically different from WT but span less than half of the possible range and the set of substitution outcomes cluster nearer WT[11; 12] than to an inactive variant. | Neutral score < 0.7<br>Rheostat score < 0.5<br>Toggle score < 0.67<br>(average closer to WT than to inactive) | 34 | 73 |
| RHEOSTAT | Substitution outcomes sample ≥50% of the accessible range[9] | Rheostat score > 0.5 | 125 | 101 |
| ADVERSE | Substitutions have detectable function, but most are highly detrimental; substitution outcomes sample less than half of the possible range and cluster near the inactive variant. This category was defined during this work. | Neutral score < 0.7<br>Rheostat score < 0.5<br>Toggle score < 0.67<br>(average closer to inactive than to WT) | 29 | 9 |
| TOGGLE | >64% of substitutions at position lack detectable activity[13] | Toggle score > 0.64 | 3 | 73 |

**Supplementary figure 4. Representative histograms for Mpro positions in the six different substitution classes, using data from (a-c, e-f) the 2022 growth assay data[1] and (d) the 2024 FRET assay data[2].** The bin containing catalytically inactive Mpro (C145A) is shown by the magenta circle and the bin containing WT is shown by the green circle; in some plots, the green circle obscures the single WT variant in the WT bin. The error range determined from WT replicates is indicated by the horizontal green bar for plots from the 2022 growth study; the WT error of the 2024 FRET study was smaller than the WT bin width. **(a)** Neutral position (position 1); **(b)** Rheostat position (position 2), the substitution outcomes at this position used to create this plot are shown in **Supplementary figure 1**; **(c)** Toggle position (position 290); **(d)** Enhancing position (position 151, from 2024 FRET assay[2]); **(e)** Moderate rheostat position (position 15); **(f)** Adverse position (position 11). The relevant scores used to classify these positions are noted on each plot. The score thresholds used to make these classifications are listed in **Supplementary table 1**.

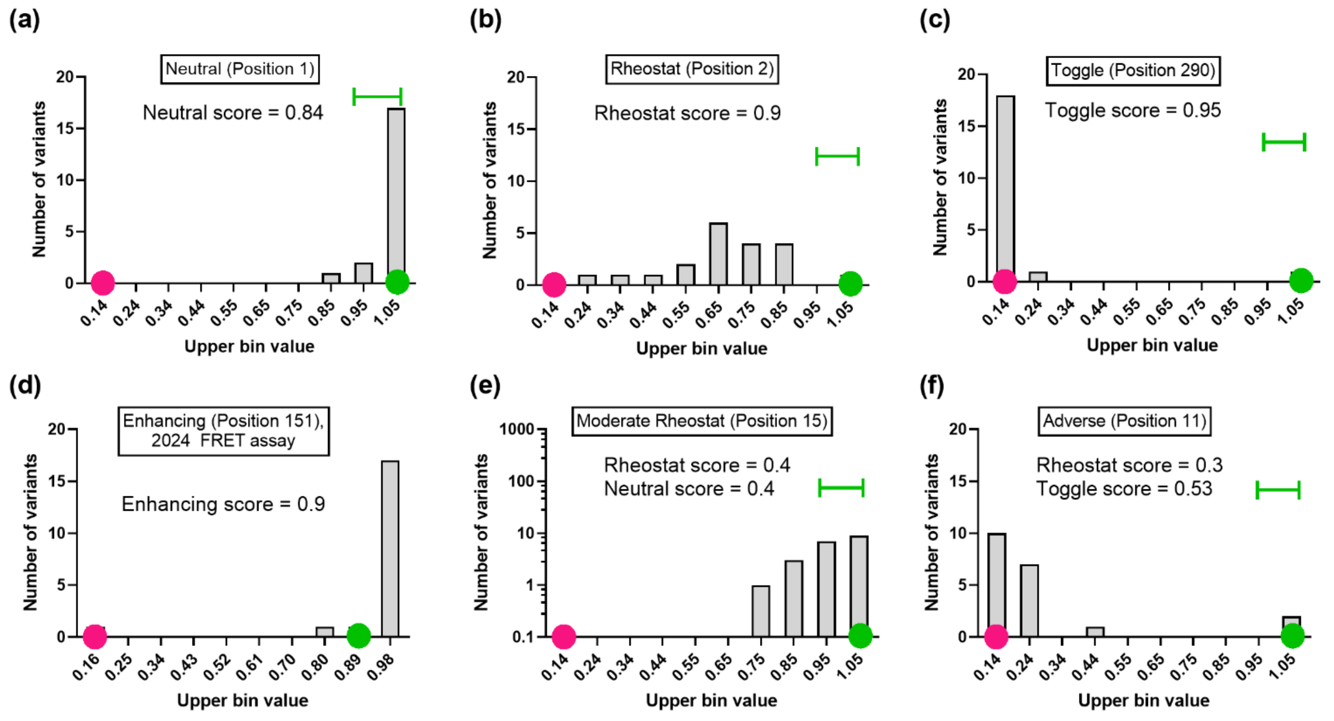

**Supplementary figure 5. Comparison of each position's DMS neutral scores (2022 growth assay[1]) with "dynamic flexibility index" (DFI) computed for Mpro[14]**. The RheoScale threshold for positional neutrality (0.7) is indicated by the vertical dotted line; scores below this threshold indicate non-neutral positions. The positions tested in this study are shown in large gold circles labelled with their respective position numbers; one of the reasons these positions were chosen is the differences in their predicted flexibilities.

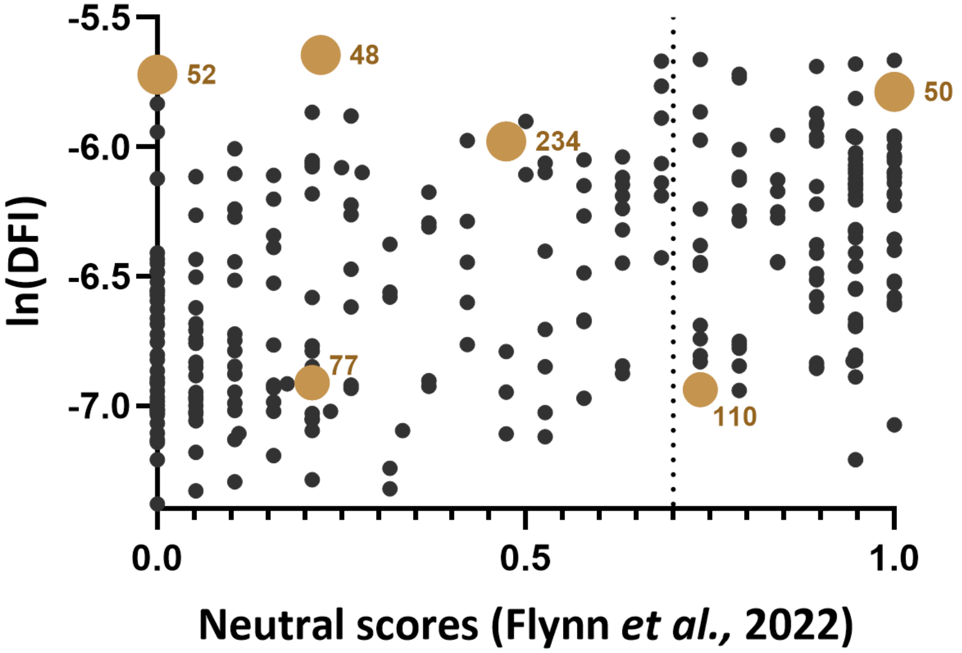

**Supplementary figure 6. Experimental design for separately measuring effects on function and *in vivo* abundance in this work.** The oval represents modified HEK293 cells that stably expressed a luciferase reporter gene integrated into its genome. These cells were transiently transfected with the reporter plasmid containing the coding region for Mpro variants. Two independent measurements were made for the WT and each Mpro variant tested in this study: (i) the signal from the luciferase assay measured "combined" effects on function and abundance changes, (ii) the indirect sandwich ELISA targeting Mpro measured *in vivo* abundance changes. The outcomes from the luciferase assays were normalized to the ELISA measurements to ascertain the effects of single substitutions on Mpro function ("Mpro outcomes" 1-3 in Methods).

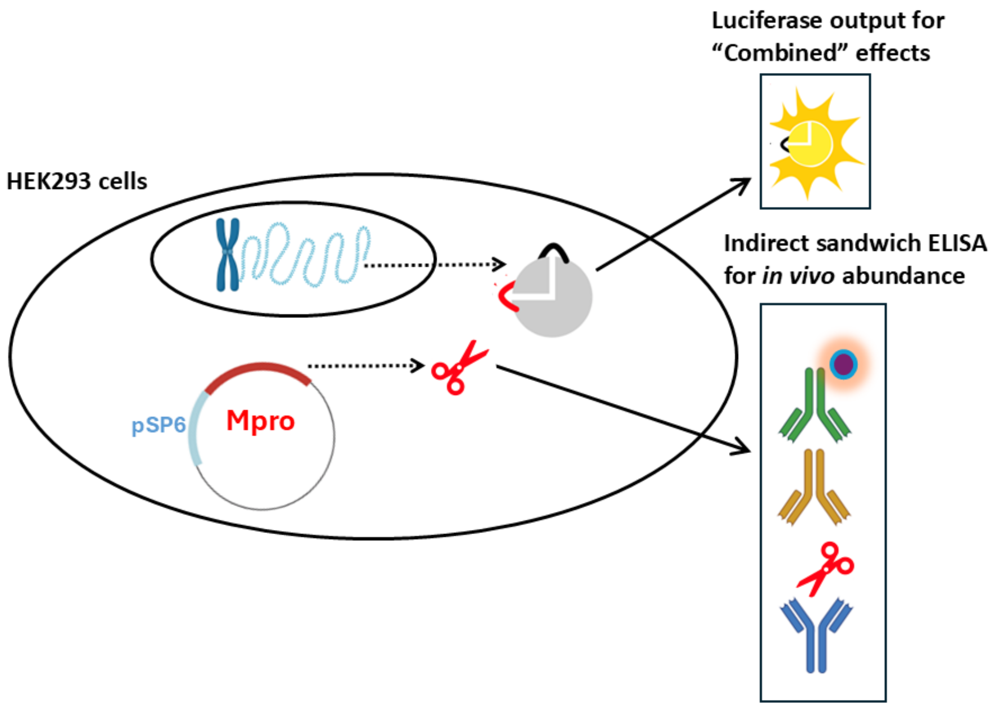

**Supplementary figure 7. Effects of single amino acid substitutions on Mpro's activity in the luciferase assay, *in vivo* abundance, and normalized function (next page).** For each of the six tested Mpro positions, ten random substitutions were made. Changes in the luciferase output, which are "combined effects", were measured (first column). Changes in *in vivo* abundance were separately assessed using an indirect sandwich ELISA (second column). These two measurements were then used to compute the abundance-normalized function for each substitution (third column; "Mpro outcome 3" in Methods). Values for Mpro WT and inactive catalysis/zero abundance were measured and all values were normalized to a WT value of 1.

These three outcomes are shown in one row for each position; note that the X-axes for the three plots are ordered the same. **(a-c)** D48, **(d-f)** L50, **(g-i)** P52, **(j-l)** V77, **(m-o)** Q110, **(p-r)** A234. On each plot, the green regions delineate the standard deviation of all WT values measured with the six sets of substitutions. For each Mpro substitution, 4-6 technical replicates for two biological replicates were measured (open circles); bars show the average for each variant and error bars are standard deviations. Error propagation for (c) is described in Methods. Variants with outcomes statistically different from WT were identified using one-way ANOVAs with Dunnett's correction (****$p < 0.0001$, ***$p<0.001$, **$p<0.01$, *$p<0.1$); significance asterisks are shown on the X-axis labels. Average values for all variants are available in the **Supplementary data**.

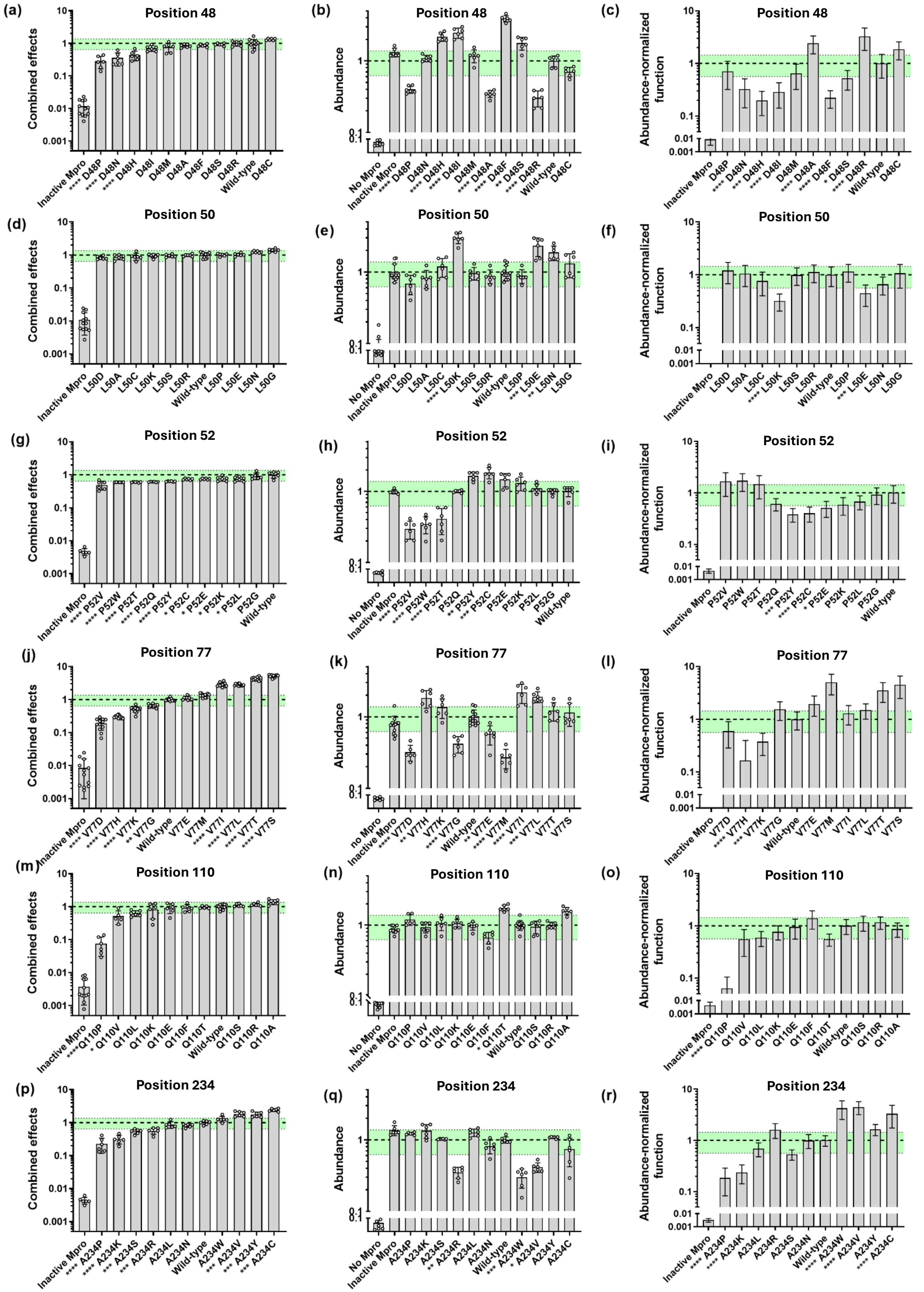
(a)
Position 48
Combined effects
Inactive Mpro, **** D48P, **** D48N, **** D48H, D48I, D48M, D48A, D48F, D48S, D48R, Wild-type, D48C
(b)
Position 48
Abundance
No Mpro, Inactive Mpro, **** D48P, D48N, **** D48H, **** D48I, D48M, **** D48A, **** D48F, ** D48S, **** D48R, Wild-type, D48C
(c)
Position 48
Abundance-normalized function
Inactive Mpro, D48P, **** D48N, ** D48H, **** D48I, D48M, **** D48A, **** D48F, * D48S, **** D48R, Wild-type, D48C
(d)
Position 50
Combined effects
Inactive Mpro, L50D, L50A, L50C, L50K, L50S, L50R, Wild-type, L50P, L50E, L50N, L50G
(e)
Position 50
Abundance
No Mpro, Inactive Mpro, L50D, L50A, L50C, **** L50K, L50S, L50R, Wild-type, L50P, *** L50E, ** L50N, L50G
(f)
Position 50
Abundance-normalized function
Inactive Mpro, L50D, L50A, L50C, **** L50K, L50S, L50R, Wild-type, L50P, *** L50E, L50N, L50G
(g)
Position 52
Combined effects
Inactive Mpro, **** P52V, **** P52W, **** P52T, **** P52Q, **** P52Y, * P52C, * P52E, * P52K, * P52L, P52G, Wild-type
(h)
Position 52
Abundance
No Mpro, Inactive Mpro, **** P52V, **** P52W, **** P52T, P52Q, ** P52Y, *** P52C, P52E, P52K, P52L, P52G, Wild-type
(i)
Position 52
Abundance-normalized function
Inactive Mpro, P52V, P52W, P52T, P52Q, *** P52Y, **** P52C, * P52E, P52K, P52L, P52G, Wild-type
(j)
Position 77
Combined effects
Inactive Mpro, **** V77D, **** V77H, **** V77K, ** V77G, Wild-type, V77E, V77M, **** V77I, **** V77L, **** V77T, **** V77S
(k)
Position 77
Abundance
no Mpro, Inactive Mpro, **** V77D, ** V77H, V77K, **** V77G, Wild-type, ** V77E, **** V77M, **** V77I, *** V77L, V77T, V77S
(l)
Position 77
Abundance-normalized function
Inactive Mpro, V77D, **** V77H, *** V77K, V77G, Wild-type, V77E, V77M, V77I, V77L, V77T, V77S
(m)
Position 110
Combined effects
Inactive Mpro, ****Q110P, * Q110V, Q110L, Q110K, Q110E, Q110F, Q110T, Wild-type, Q110S, Q110R, Q110A
(n)
Position 110
Abundance
No Mpro, Inactive Mpro, Q110P, Q110V, Q110L, Q110K, Q110E, Q110F, * Q110T, Wild-type, Q110S, Q110R, Q110A
(o)
Position 110
Abundance-normalized function
Inactive Mpro, **** Q110P, Q110V, Q110L, Q110K, Q110E, Q110F, Q110T, Wild-type, Q110S, Q110R, Q110A
(p)
Position 234
Combined effects
Inactive Mpro, **** A234P, **** A234K, *** A234S, *** A234R, A234L, A234N, Wild-type, A234W, *** A234V, *** A234Y, *** A234C
(q)
Position 234
Abundance
No Mpro, Inactive Mpro, A234P, A234K, A234S, ** A234R, A234L, A234N, Wild-type, *** A234W, * A234V, A234Y, A234C
(r)
Position 234
Abundance-normalized function
Inactive Mpro, **** A234P, **** A234K, A234L, A234R, A234S, A234N, Wild-type, **** A234W, **** A234V, A234Y, **** A234C

**Supplementary figure 8. Comparison of substitution outcomes from the two published DMS studies with the three experimental measurements in the current study.** Each panel compares values from the DMS datasets – **(a, c, e)** 2022 growth assay[1] and **(b, d, f)** the 2024 FRET assay[2] – with the log-transformed parameters measured in this study: **(a, b)** luciferase output (combined effects), **(c, d)** *in vivo* abundance, and **(e, f)** abundance-normalized function. The dotted vertical and horizontal lines correspond to the average WT values of each assay. The strongest correlations were observed for luciferase output, suggesting that our combined-effect measurement is the most directly comparable to the DMS growth and FRET assays. This is expected, given that both DMS assays report a composite phenotype that reflects contributions from both function and abundance, similar to our luciferase output.

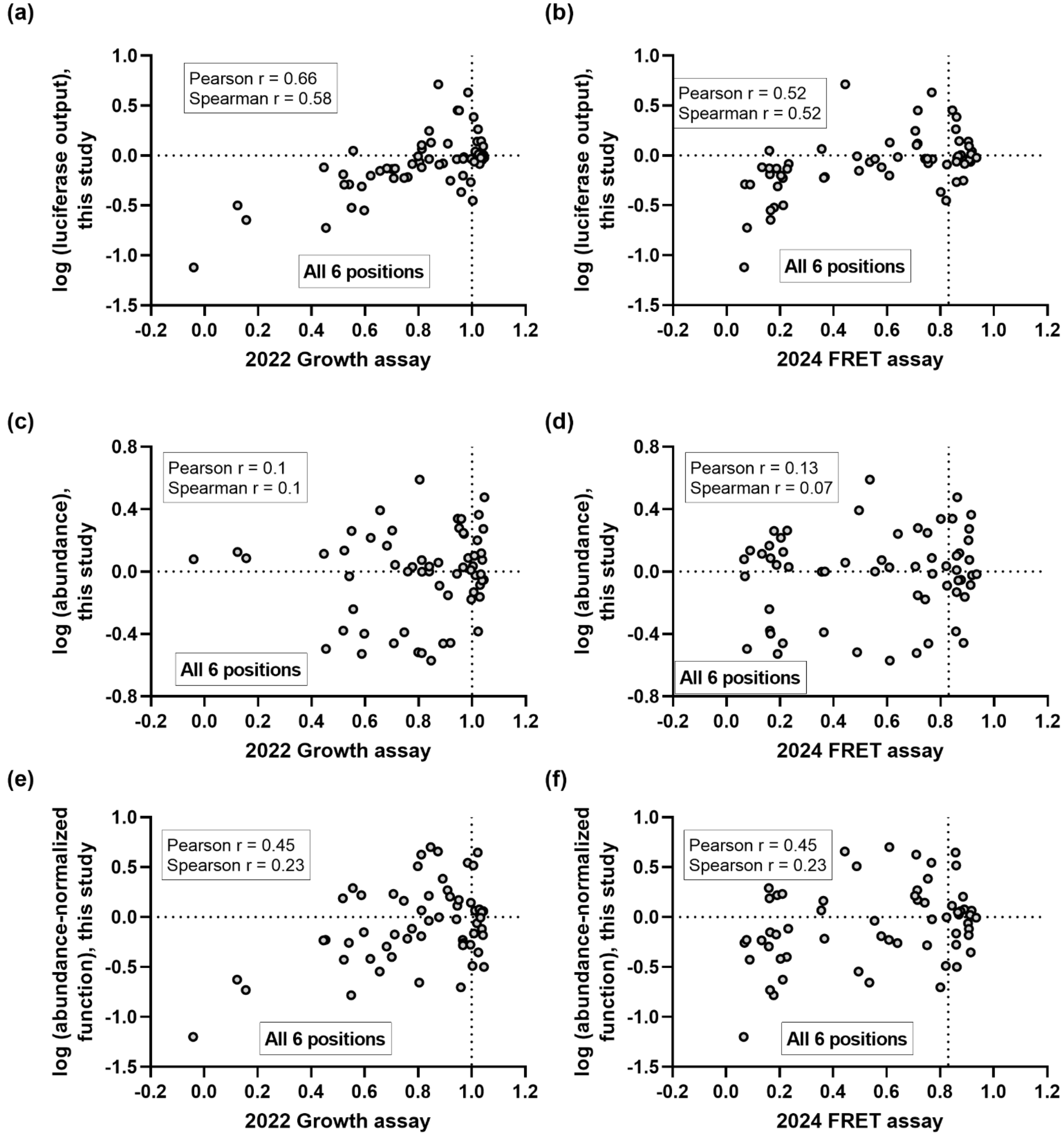

**Supplementary figure 9. Optimization of Mpro plasmid transfection in the luciferase assay.** Cells were transfected with increasing amounts of wild-type Mpro plasmid to determine the linear dynamic range of the assay. Each condition also included co-transfection with a constitutively expressed *Renilla* luciferase plasmid as an internal transfection control. The y-axis represents the ratio of firefly luciferase signal (cleaved by Mpro) to *Renilla* luciferase signal, plotted on a logarithmic scale. Luciferase signal increased with Mpro amounts up to ~60 ng, after which signal plateaued or declined, likely due to cytotoxicity. Based on this titration, 15 ng of Mpro plasmid was selected for experimental assays as it lies within the linear, non-saturating range. The open dots represent individual measurements, the bars show the average luciferase output from six technical replicates, and the error bar corresponds to the standard deviation of the average. This plot shows that the assay is sensitive to changes in Mpro abundance.

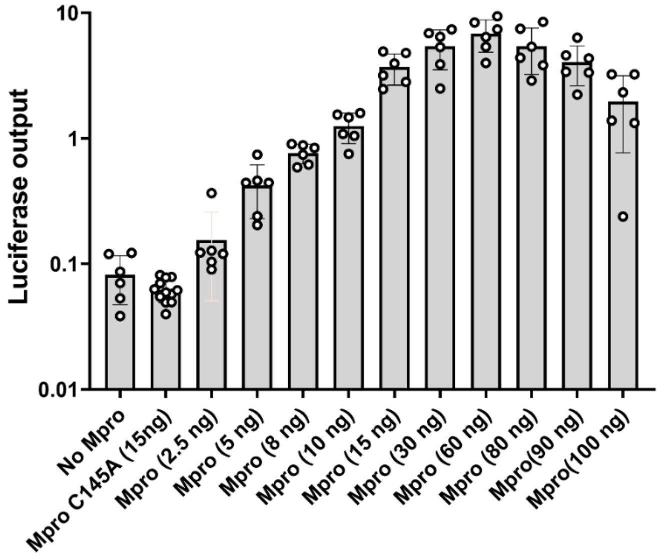

**Supplementary figure 10. Plots of abundance-normalized function versus luciferase output that arise from combined effects on abundance and activity**. **(a)** D48, **(b)** L50, **(c)** P52, **(d)** V77, **(e)** Q110, **(f)** A234. These plots illustrate that changes in abundance make varied contributions to the overall "combined" luciferase output (i) among the six positions tested and (ii) within the set of substitutions for each position. Each data point corresponds to the average values shown in **Supplementary figure 7** and listed in **Supplementary data**; error bars correspond to the standard deviations for each measurement; deviations from the diagonal line indicate substitutions for which changes in abundance make significant contributions to the un-normalized luciferase outcome.

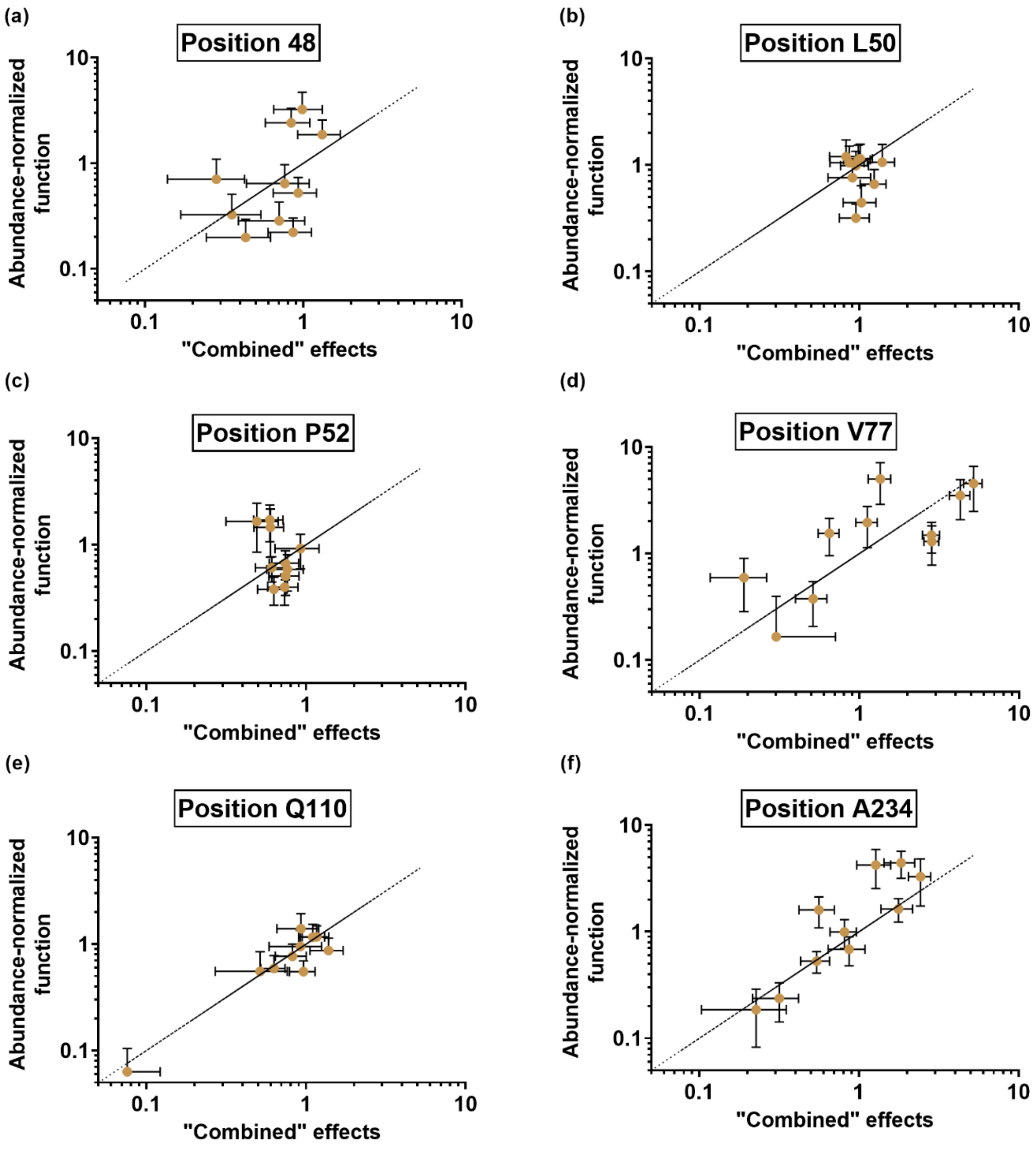

**Supplementary figure 11. Substitution effects on abundance and abundance-normalized function show a negative relationship at several positions. (a)** D48, **(b)** L50, **(c)** P52, **(d)** V77, **(e)** Q110, **(f)** A234. Each data point corresponds to the average values shown in **Supplementary figure 7** and listed in the **Supplementary data**. Standard deviations for measurements are shown by black bars for both X and Y values.

Three positions (48, 50, 52) show a strong correlation, indicating a “stability-function tradeoff”[15-18]; these three positions are on or near solvent exposed region of the structure and are far from the active site (main text **Figure 5**). In contrast, position 77 does not show this tradeoff, which suggests that different protein positions can make different contributions to function and abundance. Most substitutions at positions (110 and 235) have very little effect on abundance, which means that comparisons do not return a meaningful correlation.

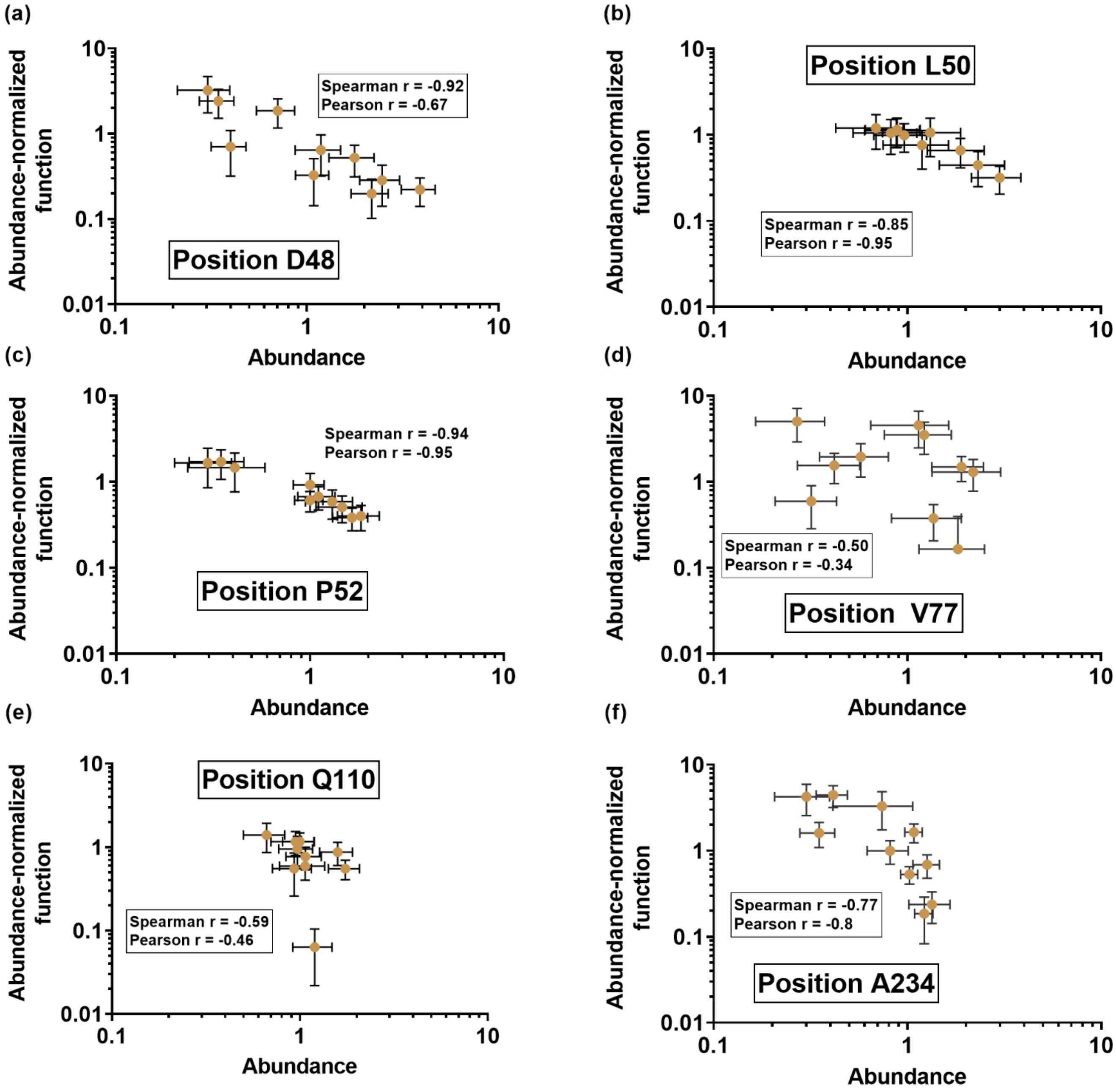

**Supplementary table 2.** Comparison of sequence and structural features of the six tested Mpro positions. Accessible surface area was measured on PDB 5R7Y[19] using USCF Chimera for the WT amino acid listed.

| Position # | Substitution sensitivity class | | | Accessible surface area[20] | Secondary structure element |
|---|---|---|---|---|---|
| | Luciferase output | Abundance | Function | | |
| D48 | Rheostat | Rheostat | Rheostat | 71.52 | Helix |
| L50 | Neutral | Moderate Rheostat | Neutral | 92.20 | Helix |
| P52 | Rheostat | Moderate Rheostat | Moderate Rheostat | 55.31 | Coil |
| V77 | Rheostat | Rheostat | Rheostat | 22.78 | Sheet |
| Q110 | Neutral | Neutral | Neutral | 68.94 | Coil |
| A234 | Rheostat | Neutral | Moderate Rheostat | 11.18 | Helix |

**Supplementary figure 12 (next page). Substitution outcomes at the six tested Mpro positions compared to secondary structure propensities.** We compared each substitution outcome to its relevant secondary structure propensity. For positions in or near helices, position-specific propensities (reported in kcal/mol) were obtained from[21]: position 48 is N2, position 50 is N4, position 52 could be a C-cap if it were to extend the helix, and position 234 is C2. For β-sheet position 77, side chain propensities (reported in kcal/mol) were obtained from [22]. Secondary structure propensities were compared to the ln-transformed experimental outcomes: luciferase output arising from combined effects on abundance and activity **(column 1)**, *in vivo* abundance **(column 2)**, and abundance-normalized function **(column 3)**. Open dots represent the ln-transformed averages of the experimental values reported in **Supplementary data** and **Supplementary figure 7**; error bars represent the transformed standard deviations. Positions classified as neutral for a given outcome were excluded (empty boxes). Pearson and Spearman correlation coefficients are reported on each plot. In four cases with correlation coefficients ≥0.5, the slope, p-value and $R^2$ values of the best-fit line (thin solid line) suggest that the significance is low and would require 10 remaining substitutions to assess whether correlations are meaningful.

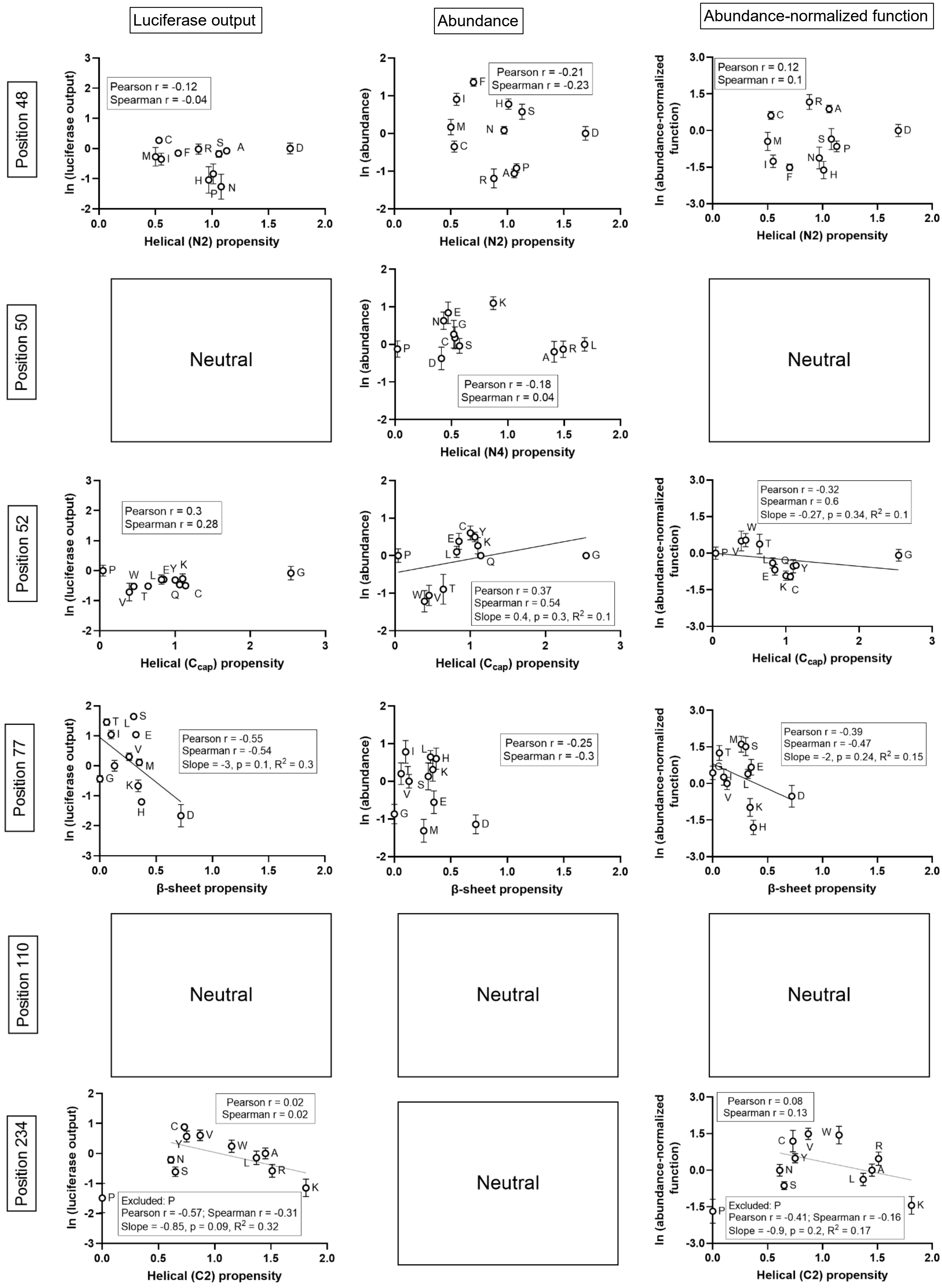

Luciferase output
Abundance
Abundance-normalized function
Position 48
Pearson r = -0.12
Spearman r = -0.04
ln (luciferase output)
Helical (N2) propensity
Pearson r = -0.21
Spearman r = -0.23
ln (abundance)
Helical (N2) propensity
Pearson r = 0.12
Spearman r = 0.1
ln (abundance-normalized function)
Helical (N2) propensity
Position 50
Neutral
ln (abundance)
Pearson r = -0.18
Spearman r = 0.04
Helical (N4) propensity
Neutral
Position 52
Pearson r = 0.3
Spearman r = 0.28
ln (luciferase output)
Helical ($C_{cap}$) propensity
ln (abundance)
Pearson r = 0.37
Spearman r = 0.54
Slope = 0.4, p = 0.3, $R^2$ = 0.1
Helical ($C_{cap}$) propensity
Pearson r = -0.32
Spearman r = 0.6
Slope = -0.27, p = 0.34, $R^2$ = 0.1
ln (abundance-normalized function)
Helical ($C_{cap}$) propensity
Position 77
ln (luciferase output)
Pearson r = -0.55
Spearman r = -0.54
Slope = -3, p = 0.1, $R^2$ = 0.3
β-sheet propensity
ln (abundance)
Pearson r = -0.25
Spearman r = -0.3
β-sheet propensity
ln (abundance-normalized function)
Pearson r = -0.39
Spearman r = -0.47
Slope = -2, p = 0.24, $R^2$ = 0.15
β-sheet propensity
Position 110
Neutral
Neutral
Neutral
Position 234
Pearson r = 0.02
Spearman r = 0.02
ln (luciferase output)
Excluded: P
Pearson r = -0.57; Spearman r = -0.31
Slope = -0.85, p = 0.09, $R^2$ = 0.32
Helical (C2) propensity
Neutral
Pearson r = 0.08
Spearman r = 0.13
ln (abundance-normalized function)
Excluded: P
Pearson r = -0.41; Spearman r = -0.16
Slope = -0.9, p = 0.2, $R^2$ = 0.17
Helical (C2) propensity

**Supplementary figure 13 (next page). Substitution outcomes at the six Mpro positions compared to amino acid side chain size.** For each of the single amino acid substitutions assessed in this study, the amino acid side chain size was obtained from solvent-accessible surface area calculations (Å$^2$) for G-X-G tripeptides as previously described[23]. Amino acid sizes were compared to the experimental outcomes: luciferase output arising from combined effects on abundance and activity **(column 1)**, *in vivo* abundance **(column 2)**, and abundance-normalized function **(column 3)**. Open dots represent the averages of the experimental values reported in **Supplementary data** and **Supplementary figure 7**; error bars represent the standard deviations. Positions classified as neutral for a given outcome were excluded (empty boxes). Pearson and Spearman correlation coefficients are reported on each plot. No meaningful correlations were observed.

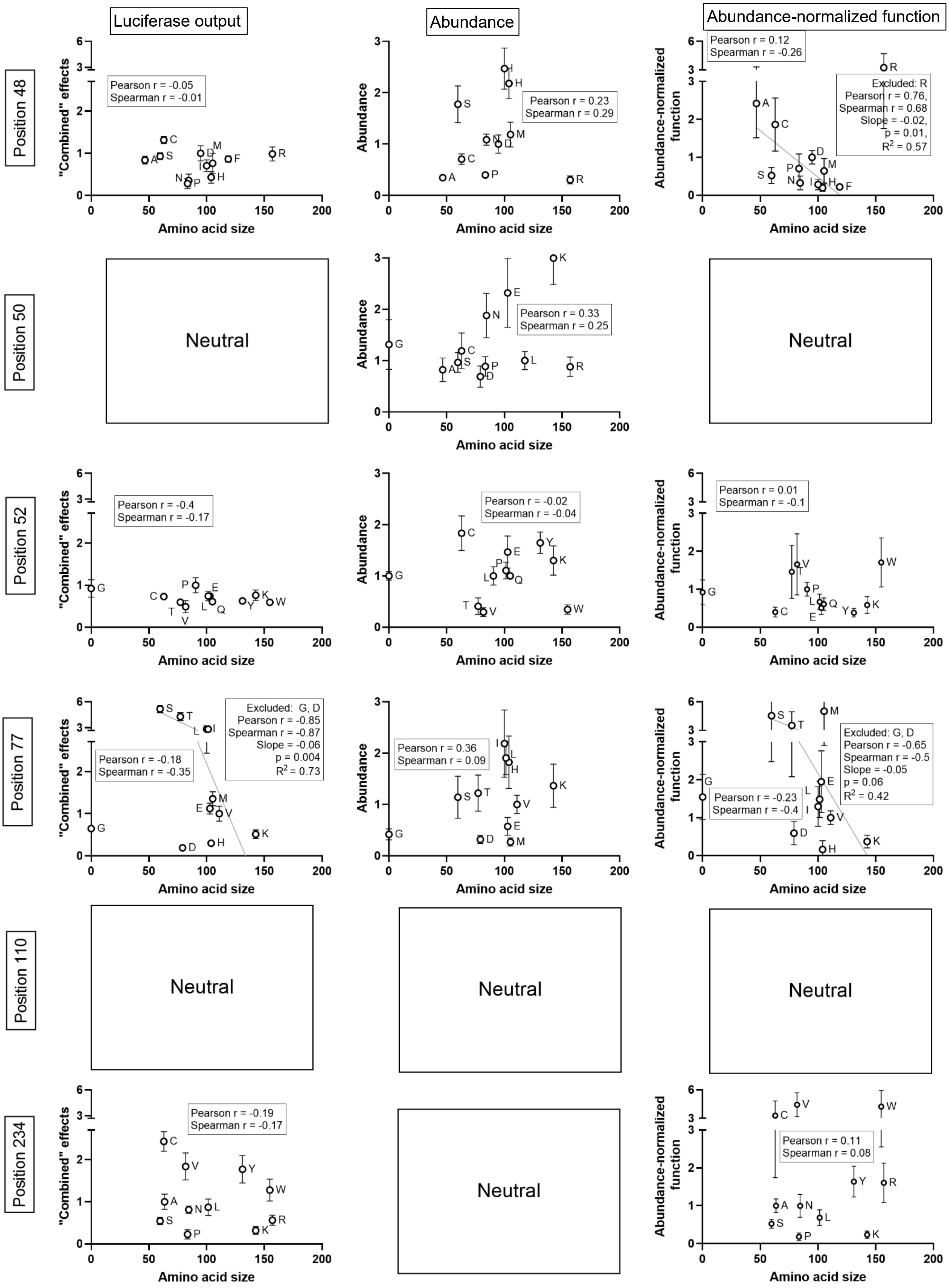

Luciferase output
Abundance
Abundance-normalized function
Position 48
Pearson r = -0.05
Spearman r = -0.01
Pearson r = 0.23
Spearman r = 0.29
Pearson r = 0.12
Spearman r = -0.26
Excluded: R
Pearson r = 0.76,
Spearman r = 0.68
Slope = -0.02,
p = 0.01,
R2 = 0.57
Position 50
Neutral
Pearson r = 0.33
Spearman r = 0.25
Neutral
Position 52
Pearson r = -0.4
Spearman r = -0.17
Pearson r = -0.02
Spearman r = -0.04
Pearson r = 0.01
Spearman r = -0.1
Position 77
Pearson r = -0.18
Spearman r = -0.35
Excluded: G, D
Pearson r = -0.85
Spearman r = -0.87
Slope = -0.06
p = 0.004
R2 = 0.73
Pearson r = 0.36
Spearman r = 0.09
Pearson r = -0.23
Spearman r = -0.4
Excluded: G, D
Pearson r = -0.65
Spearman r = -0.5
Slope = -0.05
p = 0.06
R2 = 0.42
Position 110
Neutral
Neutral
Neutral
Position 234
Pearson r = -0.19
Spearman r = -0.17
Neutral
Pearson r = 0.11
Spearman r = 0.08
"Combined" effects
Abundance
Abundance-normalized function
Amino acid size

**Supplementary figure 14. Substitution outcomes at two of six Mpro positions compared to amino acid hydrophobicity (next page).** For each of the single amino acid substitutions assessed in this study, interfacial hydrophobicity values were obtained from the Wimley–White interfacial hydrophobicity scale[24], which quantifies the free energy of partitioning amino acid side chains from water to the membrane interface (kcal/mol). These values were compared to each of the ln-transformed experimental data: luciferase output arising from combined effects on abundance and activity **(column 1)**, abundance **(column 2)**, and abundance-normalized function **(column 3)**. Open dots represent the averages of the ln-transformed experimental values reported in **Supplementary data** and **Supplementary figure 7**; error bars represent the transformed standard deviations. Positions that were neutral for the outcome compared in the column were excluded (empty boxes). The Pearson and Spearman correlation coefficients for each comparison are shown on the corresponding plots. In five cases with correlation coefficients ≥0.5, the slope, p-value and $R^2$ values of the best-fit line (thin solid lines) indicated that the relationship could be meaningful.

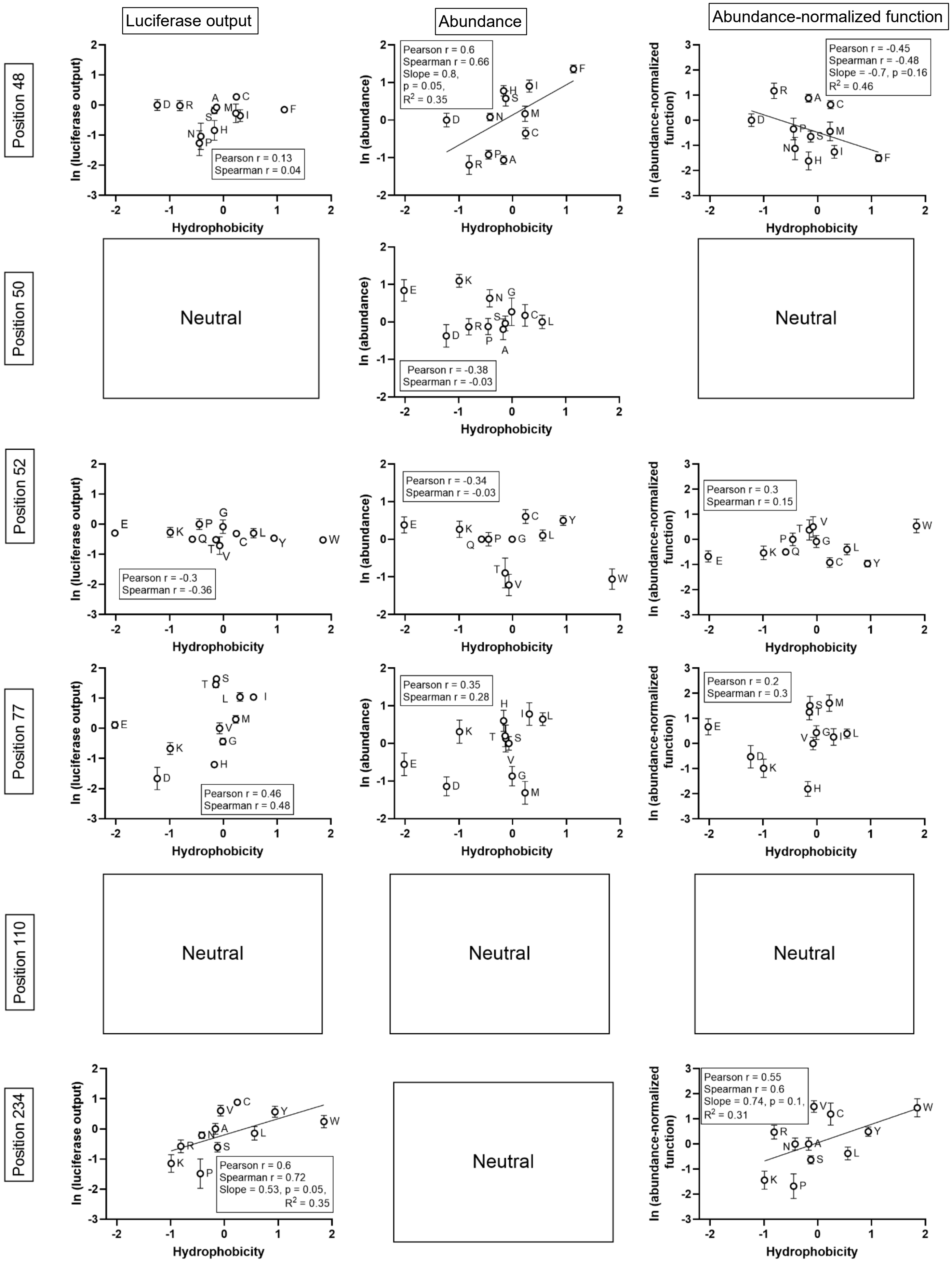

Luciferase output
Abundance
Abundance-normalized function
Position 48
Position 50
Position 52
Position 77
Position 110
Position 234
ln (luciferase output)
ln (abundance)
ln (abundance-normalized function)
Hydrophobicity
Pearson r = 0.13
Spearman r = 0.04
Pearson r = 0.6
Spearman r = 0.66
Slope = 0.8,
p = 0.05,
R2 = 0.35
Pearson r = -0.45
Spearman r = -0.48
Slope = -0.7, p =0.16
R2 = 0.46
Neutral
Pearson r = -0.38
Spearman r = -0.03
Neutral
Pearson r = -0.3
Spearman r = -0.36
Pearson r = -0.34
Spearman r = -0.03
Pearson r = 0.3
Spearman r = 0.15
Pearson r = 0.46
Spearman r = 0.48
Pearson r = 0.35
Spearman r = 0.28
Pearson r = 0.2
Spearman r = 0.3
Neutral
Neutral
Neutral
Pearson r = 0.6
Spearman r = 0.72
Slope = 0.53, p = 0.05,
R2 = 0.35
Neutral
Pearson r = 0.55
Spearman r = 0.6
Slope = 0.74, p = 0.1,
R2 = 0.31

**Supplementary table 3. Statistical analyses for experimental outcomes measured for the 60 Mpro substitutions tested in this study, relative to the WT sample measured in parallel.** As described in the main text Methods, all statistics were performed on log-transformed data.

**Supplementary table 3a.** One way ANOVA Dunnett's multiple comparisons test for **"**combined effects" from luciferase outputs.

| Dunnett's multiple comparisons test | Mean Diff. | 95.00% CI of diff. | Summary | Adjusted P Value |
|---|---|---|---|---|
| Wild-type vs. Q110A | -0.1401 | -0.4039 to 0.1236 | ns | 0.6894 |
| Wild-type vs. Q110E | 0.05379 | -0.2099 to 0.3175 | ns | 0.9992 |
| Wild-type vs. Q110F | 0.03785 | -0.2259 to 0.3016 | ns | 0.9995 |
| Wild-type vs. Q110K | 0.1366 | -0.1271 to 0.4004 | ns | 0.7172 |
| Wild-type vs. Q110L | 0.2024 | -0.06130 to 0.4662 | ns | 0.2374 |
| Wild-type vs. Q110P | 1.182 | 0.9183 to 1.446 | **** | <0.0001 |
| Wild-type vs. Q110R | -0.06995 | -0.3337 to 0.1938 | ns | 0.9929 |
| Wild-type vs. Q110S | -0.0453 | -0.3090 to 0.2184 | ns | 0.9994 |
| Wild-type vs. Q110T | 0.0126 | -0.2682 to 0.2934 | ns | 0.9999 |
| Wild-type vs. Q110V | 0.3136 | 0.03280 to 0.5944 | * | 0.0197 |
| Wild-type vs. L50A | 0.06313 | -0.1158 to 0.2420 | ns | 0.961 |
| Wild-type vs. L50C | 0.04674 | -0.1321 to 0.2256 | ns | 0.9933 |
| Wild-type vs. L50D | 0.07971 | -0.09917 to 0.2586 | ns | 0.8525 |
| Wild-type vs. L50E | -0.01726 | -0.1961 to 0.1616 | ns | 0.9997 |
| Wild-type vs. L50G | -0.1477 | -0.3266 to 0.03117 | ns | 0.1672 |
| Wild-type vs. L50K | 0.01836 | -0.1605 to 0.1972 | ns | 0.9996 |
| Wild-type vs. L50N | -0.09896 | -0.2778 to 0.07993 | ns | 0.6406 |
| Wild-type vs. L50P | -0.01052 | -0.1894 to 0.1684 | ns | 0.9998 |
| Wild-type vs. L50R | 0.002077 | -0.1768 to 0.1810 | ns | >0.9999 |
| Wild-type vs. L50S | 0.01718 | -0.1617 to 0.1961 | ns | 0.9997 |
| Wild-type vs. V77D | 0.7565 | 0.6013 to 0.9117 | **** | <0.0001 |
| Wild-type vs. V77E | -0.04853 | -0.2037 to 0.1066 | ns | 0.9745 |
| Wild-type vs. V77G | 0.1891 | 0.03396 to 0.3443 | ** | 0.0085 |
| Wild-type vs. V77H | 0.5218 | 0.3666 to 0.6770 | **** | <0.0001 |
| Wild-type vs. V77I | -0.4508 | -0.6060 to -0.2956 | **** | <0.0001 |
| Wild-type vs. V77K | 0.2977 | 0.1425 to 0.4529 | **** | <0.0001 |
| Wild-type vs. V77L | -0.4525 | -0.6077 to -0.2973 | **** | <0.0001 |
| Wild-type vs. V77M | -0.1291 | -0.2843 to 0.02607 | ns | 0.1535 |
| Wild-type vs. V77S | -0.7148 | -0.8700 to -0.5596 | **** | <0.0001 |
| Wild-type vs. V77T | -0.6316 | -0.7868 to -0.4765 | **** | <0.0001 |
| Wild-type vs. D48A | 0.06257 | -0.1403 to 0.2654 | ns | 0.9853 |
| Wild-type vs. D48C | -0.1353 | -0.3381 to 0.06754 | ns | 0.4019 |
| Wild-type vs. D48F | 0.0494 | -0.1534 to 0.2522 | ns | 0.9962 |
| Wild-type vs. D48H | 0.3661 | 0.1633 to 0.5689 | **** | <0.0001 |
| Wild-type vs. D48I | 0.1428 | -0.06005 to 0.3456 | ns | 0.335 |
| Wild-type vs. D48M | 0.1195 | -0.08339 to 0.3223 | ns | 0.563 |
| Wild-type vs. D48N | 0.4655 | 0.2627 to 0.6683 | **** | <0.0001 |
| Wild-type vs. D48P | 0.5667 | 0.3639 to 0.7695 | **** | <0.0001 |
| Wild-type vs. D48R | -0.00447 | -0.2073 to 0.1984 | ns | >0.9999 |

| Dunnett's multiple comparisons test | Mean Diff. | 95.00% CI of diff. | Summary | Adjusted P Value |
|---|---|---|---|---|
| Wild-type vs. D48S | 0.01688 | -0.1860 to 0.2197 | ns | 0.9997 |
| Wild-type vs. P52C | 0.1273 | 0.01712 to 0.2374 | * | 0.0156 |
| Wild-type vs. P52E | 0.1201 | 0.009992 to 0.2303 | * | 0.0257 |
| Wild-type vs. P52G | 0.03465 | -0.07548 to 0.1448 | ns | 0.9686 |
| Wild-type vs. P52K | 0.1137 | 0.003537 to 0.2238 | * | 0.0398 |
| Wild-type vs. P52L | 0.1252 | 0.01506 to 0.2353 | * | 0.018 |
| Wild-type vs. P52Q | 0.2072 | 0.09710 to 0.3174 | **** | <0.0001 |
| Wild-type vs. P52T | 0.2153 | 0.1052 to 0.3254 | **** | <0.0001 |
| Wild-type vs. P52V | 0.3161 | 0.2060 to 0.4262 | **** | <0.0001 |
| Wild-type vs. P52W | 0.2183 | 0.1081 to 0.3284 | **** | <0.0001 |
| Wild-type vs. P52Y | 0.1929 | 0.08280 to 0.3031 | **** | <0.0001 |
| Wild-type vs. A234C | -0.3868 | -0.5479 to -0.2257 | **** | <0.0001 |
| Wild-type vs. A234K | 0.5121 | 0.3510 to 0.6732 | **** | <0.0001 |
| Wild-type vs. A234L | 0.06716 | -0.09393 to 0.2282 | ns | 0.8505 |
| Wild-type vs. A234N | 0.0903 | -0.07079 to 0.2514 | ns | 0.5637 |
| Wild-type vs. A234P | 0.6816 | 0.5205 to 0.8427 | **** | <0.0001 |
| Wild-type vs. A234R | 0.2569 | 0.09579 to 0.4180 | *** | 0.0003 |
| Wild-type vs. A234S | 0.2669 | 0.1058 to 0.4280 | *** | 0.0002 |
| Wild-type vs. A234V | -0.2611 | -0.4222 to -0.1000 | *** | 0.0003 |
| Wild-type vs. A234W | -0.1015 | -0.2626 to 0.05962 | ns | 0.426 |
| Wild-type vs. A234Y | -0.2444 | -0.4055 to -0.08331 | *** | 0.0007 |

**Supplementary table 3b.** One way ANOVA Dunnett's multiple comparisons test for *in vivo* abundance measurements.

| Dunnett's multiple comparisons test | Mean Diff. | 95.00% CI of diff. | Summary | Adjusted P Value |
|---|---|---|---|---|
| Wild-type vs. Q110A | -0.2036 | -0.4122 to 0.005021 | ns | 0.06 |
| Wild-type vs. Q110E | 0.01162 | -0.1970 to 0.2202 | ns | 0.9998 |
| Wild-type vs. Q110F | 0.1795 | -0.02914 to 0.3881 | ns | 0.1359 |
| Wild-type vs. Q110K | -0.031 | -0.2396 to 0.1776 | ns | 0.9994 |
| Wild-type vs. Q110L | -0.02343 | -0.2320 to 0.1852 | ns | 0.9996 |
| Wild-type vs. Q110P | -0.07873 | -0.2873 to 0.1299 | ns | 0.9489 |
| Wild-type vs. Q110R | -0.00234 | -0.2109 to 0.2063 | ns | >0.9999 |
| Wild-type vs. Q110S | 0.0266 | -0.1820 to 0.2352 | ns | 0.9996 |
| Wild-type vs. Q110T | -0.2454 | -0.4540 to -0.03681 | * | 0.0116 |
| Wild-type vs. Q110V | 0.03121 | -0.1774 to 0.2398 | ns | 0.9994 |
| Wild-type vs. L50A | 0.08854 | -0.1320 to 0.3090 | ns | 0.9248 |
| Wild-type vs. L50C | -0.06942 | -0.2899 to 0.1511 | ns | 0.9868 |
| Wild-type vs. L50D | 0.1707 | -0.04979 to 0.3912 | ns | 0.2325 |
| Wild-type vs. L50E | -0.3577 | -0.5783 to -0.1372 | *** | 0.0001 |
| Wild-type vs. L50G | -0.1023 | -0.3228 to 0.1182 | ns | 0.8351 |

| Dunnett's multiple comparisons test | Mean Diff. | 95.00% CI of diff. | Summary | Adjusted P Value |
|---|---|---|---|---|
| Wild-type vs. L50K | -0.4809 | -0.7014 to -0.2604 | **** | <0.0001 |
| Wild-type vs. L50N | -0.2742 | -0.4948 to -0.05373 | ** | 0.0063 |
| Wild-type vs. L50P | 0.05112 | -0.1694 to 0.2716 | ns | 0.9991 |
| Wild-type vs. L50R | 0.05423 | -0.1663 to 0.2747 | ns | 0.999 |
| Wild-type vs. L50S | 0.01308 | -0.2074 to 0.2336 | ns | 0.9998 |
| Wild-type vs. V77D | 0.4954 | 0.3055 to 0.6853 | **** | <0.0001 |
| Wild-type vs. V77E | 0.2496 | 0.05967 to 0.4395 | ** | 0.0032 |
| Wild-type vs. V77G | 0.3793 | 0.1894 to 0.5692 | **** | <0.0001 |
| Wild-type vs. V77H | -0.2569 | -0.4468 to -0.06701 | ** | 0.0022 |
| Wild-type vs. V77I | -0.3329 | -0.5228 to -0.1429 | **** | <0.0001 |
| Wild-type vs. V77K | -0.128 | -0.3179 to 0.06188 | ns | 0.3989 |
| Wild-type vs. V77L | -0.2857 | -0.4756 to -0.09581 | *** | 0.0005 |
| Wild-type vs. V77M | 0.5745 | 0.3846 to 0.7644 | **** | <0.0001 |
| Wild-type vs. V77S | -0.0476 | -0.2375 to 0.1423 | ns | 0.999 |
| Wild-type vs. V77T | -0.08345 | -0.2734 to 0.1065 | ns | 0.8768 |
| Wild-type vs. **** D48A | 0.4579 | 0.2632 to 0.6527 | **** | <0.0001 |
| Wild-type vs. D48C | 0.1502 | -0.04459 to 0.3449 | ns | 0.2159 |
| Wild-type vs. **** D48F | -0.5936 | -0.7883 to -0.3988 | **** | <0.0001 |
| Wild-type vs. **** D48H | -0.3406 | -0.5354 to -0.1459 | **** | <0.0001 |
| Wild-type vs. **** D48I | -0.3934 | -0.5882 to -0.1987 | **** | <0.0001 |
| Wild-type vs. D48M | -0.07112 | -0.2659 to 0.1236 | ns | 0.9335 |
| Wild-type vs. D48N | -0.0399 | -0.2346 to 0.1548 | ns | 0.9991 |
| Wild-type vs. **** D48P | 0.3952 | 0.2005 to 0.5900 | **** | <0.0001 |
| Wild-type vs. **** D48R | 0.5234 | 0.3287 to 0.7182 | **** | <0.0001 |
| Wild-type vs. ** D48S | -0.2466 | -0.4413 to -0.05183 | ** | 0.006 |
| Wild-type vs. *** P52C | -0.2626 | -0.4376 to -0.08755 | *** | 0.0007 |
| Wild-type vs. P52E | -0.1625 | -0.3375 to 0.01249 | ns | 0.0821 |
| Wild-type vs. P52G | -0.00477 | -0.1798 to 0.1702 | ns | >0.9999 |
| Wild-type vs. P52K | -0.1118 | -0.2868 to 0.06323 | ns | 0.4194 |
| Wild-type vs. P52L | -0.04539 | -0.2204 to 0.1296 | ns | 0.9916 |
| Wild-type vs. P52Q | -0.00527 | -0.1803 to 0.1697 | ns | 0.9999 |
| Wild-type vs. **** P52T | 0.4142 | 0.2392 to 0.5892 | **** | <0.0001 |
| Wild-type vs. **** P52V | 0.5375 | 0.3625 to 0.7126 | **** | <0.0001 |
| Wild-type vs. **** P52W | 0.4692 | 0.2942 to 0.6443 | **** | <0.0001 |
| Wild-type vs. ** P52Y | -0.2192 | -0.3942 to -0.04420 | ** | 0.0068 |
| Wild-type vs. A234C | 0.1709 | -0.1585 to 0.5003 | ns | 0.6662 |
| Wild-type vs. A234K | -0.1191 | -0.4485 to 0.2103 | ns | 0.9374 |
| Wild-type vs. A234L | -0.1004 | -0.4298 to 0.2290 | ns | 0.9802 |
| Wild-type vs. A234N | 0.09736 | -0.2320 to 0.4268 | ns | 0.9844 |
| Wild-type vs. A234P | -0.08789 | -0.4173 to 0.2415 | ns | 0.9909 |
| Wild-type vs. ** A234R | 0.4611 | 0.1317 to 0.7905 | ** | 0.0018 |
| Wild-type vs. A234S | -0.01209 | -0.3415 to 0.3173 | ns | 0.9999 |
| Wild-type vs. * A234V | 0.3859 | 0.05653 to 0.7153 | * | 0.0133 |
| Wild-type vs. *** A234W | 0.5402 | 0.2108 to 0.8696 | *** | 0.0002 |
| Wild-type vs. A234Y | -0.03459 | -0.3640 to 0.2948 | ns | 0.9996 |

**Supplementary table 3c.** One way ANOVA Dunnett's multiple comparisons test for abundance-normalized function computed as described in Methods.

| Dunnett's multiple comparisons test | Mean Diff. | 95.00% CI of diff. | Summary | Adjusted P Value |
|---|---|---|---|---|
| Wild-type vs. Q110A | 0.1307 | -0.2812 to 0.5427 | ns | 0.9819 |
| Wild-type vs. Q110E | 0.05052 | -0.3614 to 0.4625 | ns | 0.9996 |
| Wild-type vs. Q110F | -0.3959 | -0.8078 to 0.01608 | ns | 0.0669 |
| Wild-type vs. Q110K | 0.2323 | -0.1797 to 0.6442 | ns | 0.6189 |
| Wild-type vs. Q110L | 0.4092 | -0.002704 to 0.8212 | ns | 0.0525 |
| Wild-type vs. Q110P | 0.9368 | 0.5249 to 1.349 | **** | <0.0001 |
| Wild-type vs. Q110R | -0.1677 | -0.5796 to 0.2442 | ns | 0.9087 |
| Wild-type vs. Q110S | -0.1615 | -0.5735 to 0.2504 | ns | 0.9262 |
| Wild-type vs. Q110T | 0.4487 | 0.01020 to 0.8873 | * | 0.0418 |
| Wild-type vs. Q110V | 0.4465 | 0.007962 to 0.8850 | * | 0.0435 |
| Wild-type vs. L50A | -0.05064 | -0.5600 to 0.4588 | ns | 0.9997 |
| Wild-type vs. L50C | 0.2389 | -0.2705 to 0.7483 | ns | 0.8124 |
| Wild-type vs. L50D | -0.1983 | -0.7077 to 0.3111 | ns | 0.9286 |
| Wild-type vs. L50E | 0.5557 | 0.04630 to 1.065 | *** | 0.0244 |
| Wild-type vs. L50G | -0.06092 | -0.5703 to 0.4485 | ns | 0.9996 |
| Wild-type vs. L50K | 0.6824 | 0.1730 to 1.192 | **** | 0.0027 |
| Wild-type vs. L50N | 0.3398 | -0.1696 to 0.8492 | ns | 0.4019 |
| Wild-type vs. L50P | -0.1448 | -0.6542 to 0.3646 | ns | 0.9909 |
| Wild-type vs. L50R | -0.1196 | -0.6290 to 0.3898 | ns | 0.9965 |
| Wild-type vs. L50S | 0.01265 | -0.4968 to 0.5220 | ns | >0.9999 |
| Wild-type vs. V77D | 0.4079 | -0.7539 to 1.570 | ns | 0.9466 |
| Wild-type vs. V77E | -0.9491 | -2.111 to 0.2127 | ns | 0.168 |
| Wild-type vs. V77G | -0.5446 | -1.706 to 0.6172 | ns | 0.7713 |
| Wild-type vs. V77H | 0.835 | -0.3268 to 1.997 | ns | 0.2889 |
| Wild-type vs. V77I | -0.2975 | -1.459 to 0.8643 | ns | 0.9919 |
| Wild-type vs. V77K | 0.6252 | -0.5366 to 1.787 | ns | 0.6277 |
| Wild-type vs. V77L | -0.485 | -1.647 to 0.6768 | ns | 0.8627 |
| Wild-type vs. V77M | -4.012 | -5.174 to -2.851 | **** | <0.0001 |
| Wild-type vs. V77S | -3.536 | -4.698 to -2.374 | **** | <0.0001 |
| Wild-type vs. V77T | -2.507 | -3.669 to -1.345 | **** | <0.0001 |
| Wild-type vs. D48A | -1.419 | -2.207 to -0.6300 | **** | <0.0001 |
| Wild-type vs. D48C | -0.8644 | -1.653 to -0.07585 | * | 0.0234 |
| Wild-type vs. D48F | 0.7787 | -0.009806 to 1.567 | ns | 0.0549 |
| Wild-type vs. D48H | 0.8021 | 0.01358 to 1.591 | * | 0.0439 |
| Wild-type vs. D48I | 0.7153 | -0.07318 to 1.504 | ns | 0.0979 |
| Wild-type vs. D48M | 0.3565 | -0.4320 to 1.145 | ns | 0.8419 |
| Wild-type vs. D48N | 0.6746 | -0.1139 to 1.463 | ns | 0.1383 |
| Wild-type vs. D48P | 0.2944 | -0.4941 to 1.083 | ns | 0.9445 |
| Wild-type vs. D48R | -2.233 | -3.022 to -1.445 | **** | <0.0001 |
| Wild-type vs. D48S | 0.4776 | -0.3109 to 1.266 | ns | 0.5269 |
| Wild-type vs. P52C | 0.6006 | -0.06150 to 1.263 | ns | 0.0937 |
| Wild-type vs. P52E | 0.4922 | -0.1699 to 1.154 | ns | 0.2458 |
| Wild-type vs. P52G | 0.07892 | -0.5832 to 0.7410 | ns | 0.9995 |

| Dunnett's multiple comparisons test | Mean Diff. | 95.00% CI of diff. | Summary | Adjusted P Value |
|---|---|---|---|---|
| Wild-type vs. P52K | 0.4142 | -0.2479 to 1.076 | ns | 0.4341 |
| Wild-type vs. P52L | 0.3287 | -0.3334 to 0.9908 | ns | 0.6987 |
| Wild-type vs. P52Q | 0.3916 | -0.2706 to 1.054 | ns | 0.5005 |
| Wild-type vs. P52T | -0.46 | -1.122 to 0.2021 | ns | 0.3152 |
| Wild-type vs. P52V | -0.6557 | -1.318 to 0.006416 | ns | 0.0536 |
| Wild-type vs. P52W | -0.7091 | -1.371 to -0.04696 | * | 0.0299 |
| Wild-type vs. P52Y | 0.6182 | -0.04389 to 1.280 | ns | 0.0787 |
| Wild-type vs. A234C | -2.288 | -3.577 to -0.9994 | **** | <0.0001 |
| Wild-type vs. A234K | 0.7631 | -0.5257 to 2.052 | ns | 0.499 |
| Wild-type vs. A234L | 0.3138 | -0.9750 to 1.603 | ns | 0.9926 |
| Wild-type vs. A234N | 0.003007 | -1.286 to 1.292 | ns | >0.9999 |
| Wild-type vs. A234P | 0.8143 | -0.4745 to 2.103 | ns | 0.4224 |
| Wild-type vs. A234R | -0.6057 | -1.894 to 0.6831 | ns | 0.7526 |
| Wild-type vs. A234S | 0.4698 | -0.8190 to 1.759 | ns | 0.9241 |
| Wild-type vs. A234V | -3.438 | -4.727 to -2.150 | **** | <0.0001 |
| Wild-type vs. A234W | -3.235 | -4.524 to -1.946 | **** | <0.0001 |
| Wild-type vs. A234Y | -0.6378 | -1.927 to 0.6510 | ns | 0.7019 |
